# Sequence context and methylation interact to shape germline mutation rate variation at CpG sites

**DOI:** 10.1101/2025.11.13.688199

**Authors:** Sheel Chandra, Ziyue Gao

## Abstract

A prominent example of sequence context-dependent mutation rate variation is the elevated transition rate at CpG sites, which is largely attributed to cytosine methylation. CpGs with different flanking sequences also exhibit mutation rate variation, but this variation is only partially correlated with context-specific methylation level. Here, we quantify the CpG mutation rate and mutagenic effect of methylation across sequence contexts. Using a regression framework that accounts for recurrent mutations, we analyze human polymorphisms from the gnomAD dataset to estimate mutation rates of unmethylated and methylated CpGs separately in each unique 4-mer or 6-mer context. We find that CpG mutation rate variation in the human genome is shaped by methylation at the focal cytosine, the flanking nucleotides, and interactions between them, suggesting distinct context-dependent mutation patterns for unmethylated and methylated cytosines. Our analysis further reveals that the context effects are driven by largely independent effects of upstream and downstream sequences. Notably, an upstream adenine markedly increases CpG mutation rates regardless of methylation status or downstream sequences. Furthermore, upstream and downstream sequences have similar effects in chimpanzee and rhesus macaque, indicating that some conserved, intrinsic sequence features shape CpG mutability. On the other hand, some inter-species differences, which are especially pronounced at methylated sites on the chimpanzee lineage, point to recent evolutionary changes, possibly in context-specificity of proteins governing DNA demethylation and repair processes.

**Author Summary:** The DNA sequence surrounding a nucleotide strongly influences how likely it is to mutate. An extreme example is the CpG dinucleotide: cytosines in CpGs mutate far more frequently than other sites in the human genome. This is related to DNA methylation, a chemical modification that occurs almost exclusively at CpGs in vertebrates and makes cytosines more prone to mutations. However, CpGs in different sequence contexts also vary in their mutation rates, and methylation level alone cannot explain this variation. To gain insight into what processes drive this variation, we estimate mutation rates for methylated and unmethylated CpGs in different sequence contexts using human genetic variation data. We find that methylation and neighboring bases interact to influence CpG mutation rates, and that the DNA sequence on either side of the CpG exerts largely independent effects. Extending our analysis to other primates reveals both conserved and species-specific patterns, with differences being especially pronounced at methylated sites on the chimpanzee lineage. Together, our results suggest that while intrinsic DNA sequence features underlie some conserved context effects on CpG mutation rate, inter-species differences may reflect recent evolutionary changes in the mechanisms that regulate DNA demethylation and repair.

## Introduction

Mutation rate varies across the human genome, even at the scale of individual nucleotides [1]. Quantifying and accounting for this mutation rate variation is important for making inferences about natural selection [2] and identifying functional coding [3–6] and non-coding regions [7]. Among the many predictors of mutation rate, one of the strongest is the local sequence context—the nucleotides flanking the mutated site [8,9]. Although context effects have been attributed to several mechanisms, including local DNA shape [10,11], biases in DNA polymerase fidelity [12], and differential repair efficiency across sequence environments [13,14], the relative importance of these mechanisms and whether they fully account for observed patterns remain open questions. Addressing these questions requires systematically characterizing how local sequence shapes site-level mutation rate variation, which can help disentangle these contributions and shed light on the mechanistic basis of mutagenesis.

Among sequence contexts, the CpG dinucleotide is both exceptionally mutable and mechanistically informative. Across vertebrates, cytosines at CpG sites undergo C>T transition at rates an order of magnitude higher than those at other genomic sites [1,15,16]. This hypermutability is directly linked to DNA methylation, as supported by two observations: first, in vertebrates, cytosine methylation occurs almost exclusively at CpG sites [17], and second, in humans, de novo C>T mutation rates at CpGs correlate strongly with germline methylation levels [18–20].

The mutagenic effect of methylation is commonly attributed to the spontaneous deamination of 5-methylcytosine (5mC). This process is distinguished from the deamination of unmethylated cytosine in two critical ways: an accelerated reaction rate [21] and a distinct chemical product. Specifically, while deamination of unmethylated cytosine yields uracil—a non-canonical base efficiently recognized and removed by DNA repair machinery [22]—deamination of 5mC produces thymine. The resulting T:G mismatch can then be converted to a T:A base pair if the mismatch is mis-repaired or left unrepaired until DNA replication [23].

However, not all C>T mutations at CpG sites arise from faulty/inefficient repair following deamination; replication-associated errors independent of deamination likely also contribute [12,24,25]. For example, analysis of de novo CpG>TpG mutations reveals strong asymmetry with regard to replication direction, showing up to 33% higher mutation rate on the lagging strand [24]. In addition, recent work on human DNA polymerase E suggests that this enzyme exhibits a bias toward mis-incorporating adenine opposite 5mC and even unmethylated cytosine in a CpG context [12,25].

CpG mutation rates vary substantially across sequence contexts [8,26]. Aggarwala and Voight [8] reported that although sperm methylation levels correlate significantly with CpG>TpG polymorphism rates in 7-mer contexts, the correlation is modest (R^2^ = 0.33), suggesting that variation in methylation level alone cannot fully explain CpG mutation rate variation across sequence contexts. A key question, then, is how the local sequence environment influences these rates. Potential mechanisms include intrinsic DNA conformational features that shape the rate of spontaneous deamination [27], the interplay between local sequence and repair enzymes in resolving T:G mismatches [14,28,29], or sequence-dependent polymerase error profiles [12,25]. In addition, context-specific DNA-binding proteins such as transcription factors can also modulate mutation rate by interfering with repair machinery [30–32]. Because 5mC and unmethylated cytosines possess distinct biochemical properties, quantifying their mutational profiles separately can help distinguish between these competing mechanisms. This necessitates a framework capable of decoupling the influence of methylation from sequence context effects, allowing for the estimation of mutation patterns at both methylated and unmethylated CpGs.

Here, we develop a modeling framework to investigate the biological determinants of CpG mutability by explicitly considering both methylation state and sequence context at each CpG site. We build on existing context-dependent mutation models by incorporating methylation as a continuous predictor, enabling us to disentangle the contributions of methylation and local sequence context while leveraging information from CpG sites across varying methylation levels. Applying this approach to human polymorphism data from the Genome Aggregation Database (gnomAD v4.0), we estimate mutation rates for both unmethylated and methylated CpGs in all unique 4-mer and 6-mer contexts. We also extend our analysis to two additional primate species to identify conserved and divergent patterns in CpG mutation rate variation. Overall, our results demonstrate that unmethylated and methylated cytosines represent fundamentally different mutational substrates with distinct context dependencies that together shape CpG mutation landscape across the genome.

## Results

### Interaction between methyl group and flanking nucleotides on CpG mutation rate

We hypothesized that the variation in CpG mutation rates across contexts is shaped by both differences in baseline mutation rates of unmethylated cytosines and differential mutagenic effects of methylation (i.e. the difference in mutation rate between methylated and unmethylated cytosines). We started by calculating the CpG transition polymorphism rate for each tetranucleotide (4-mer) context at varying methylation levels, using single nucleotide polymorphisms (SNPs) as a proxy for mutation events in the ancestors of the sampled genomes. For primary analysis, we used SNPs in the intergenic regions of the human genome from gnomAD [36], and polarized SNPs based on minor allele frequency (Methods); SNPs from the 1000 Genomes (1KG) Project were used for replication [37].

To approximate methylation levels in the male germline, we used bisulfite sequencing from human sperm [38]. We recognize that germline mutations can arise in both the male and female germlines, as well as during early embryonic development, so the most relevant methylation metric should be a weighted average of methylation levels across all these stages. However, we chose to use sperm bisulfite sequencing data as a proxy for germline methylation level for two reasons. First, the site-level methylation intensity in sperm shows the strongest correlation with CpG mutation rate and has much greater predictive power than data from any other developmental stages [39]. Second, whole-genome bisulfite sequencing data for tissues other than testis and/or sperm are much more limited and typically not corrected for genotypes of the assayed individuals, which creates biases in methylation level measurement (see Methods).

As expected, the polymorphism rate increases with germline methylation level for all CpG sites in aggregation and within each 4-mer context (Fig 1A; S1A Fig). However, this increase is approximately linear only up to ∼25% methylation, after which the curve becomes flatter. This non-linear relationship between polymorphism rate and methylation is likely driven by mutation saturation of highly mutable sites, where recurrent mutations become frequent in large cohorts [18,40]. This observation is concordant with prior studies that, at the current sample size of human genetic variation datasets, the observed polymorphism probability of CpG sites no longer scales proportionately with the underlying mutation rate [18,33,41,42].

**Fig 1.**
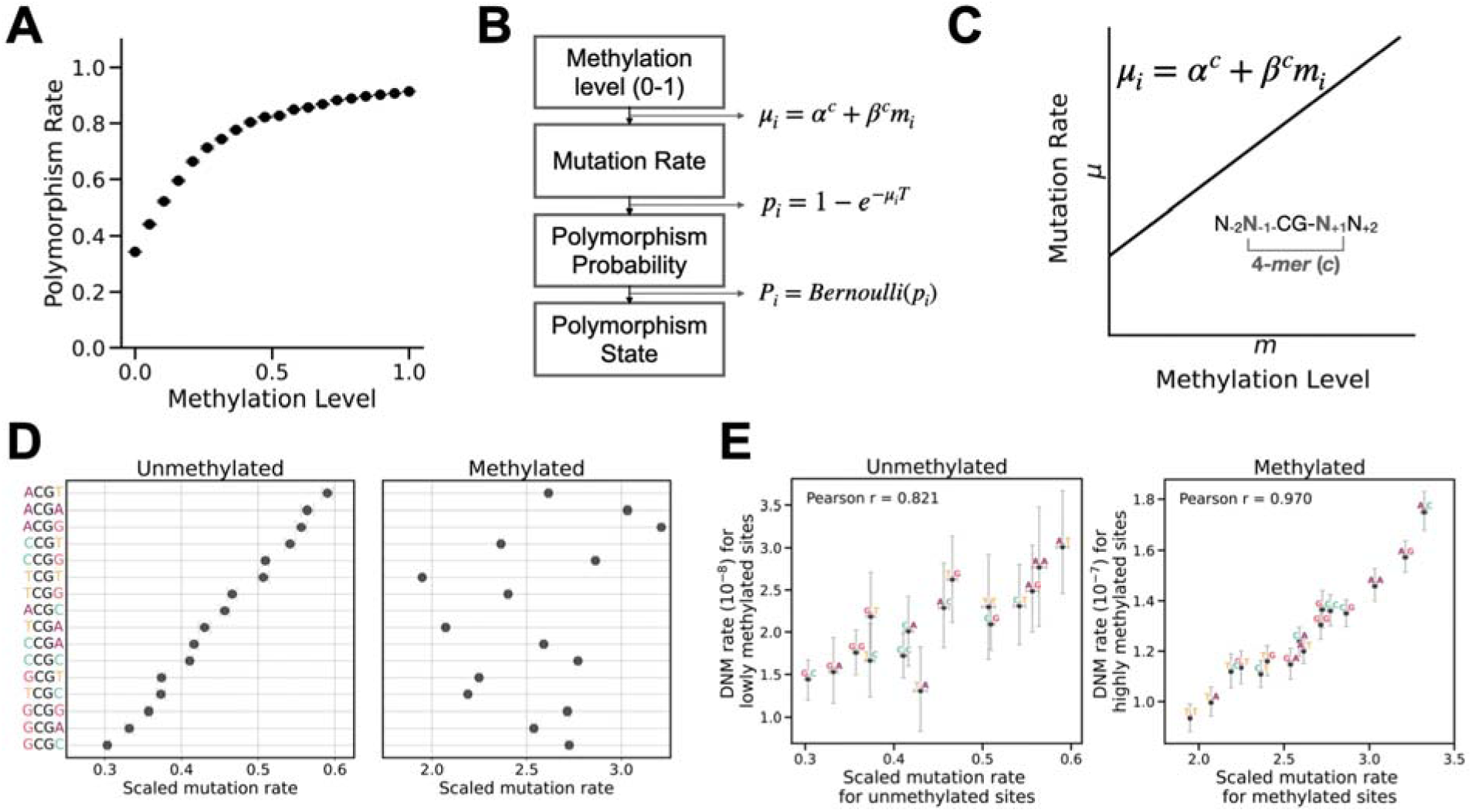
Interaction between sequence context and methylation shapes mutation rate variation at CpG sites. **A.** Polymorphism rate of CpG sites as a function of methylation levels. CpG sites were partitioned into 20 equal-width bins (ranging from [0, 0.05) to [0.95, 1]), with the polymorphism rate calculated as the proportion of sites harboring SNPs within each bin. Error bars represent 95% confidence intervals assuming binomial sampling. **B.** Proposed mutation rate model where the mutation rate for a given cytosine in a CpG within a 4-mer sequence context is a linear function of its methylation level. is the context-specific baseline rate when methylation is zero, and captures the increase in mutation rate from unmethylated to fully methylated. **C.** Schematic showing how mutation rates are related with the observed methylation level and polymorphism state at every site. **D.** Scaled mutation rate estimates for unmethylated and methylated CpGs in each 4-mer context. Contexts are sorted by descending mutation rate when unmethylated. Error bars represent 95% confidence intervals for the estimates. **E.** Comparison between scaled mutation rates of unmethylated and methylated CpGs estimated from the gnomAD dataset by our model and estimated *de novo* mutation (DNM) rates for lowly methylated (<10% methylation in sperm) and highly methylated sites (>90% methylation), respectively. 95% confidence intervals for DNM rates are estimated assuming binomial sampling.

To account for recurrent mutations, we modeled the relationship between the polymorphism probability *p* and the underlying per-generation mutation rate *μ* using an exponential transformation *p* =1 - *e*^-*µT*^, where T represents the total branch length of the coalescent tree connecting all sampled genomes. This approach adjusts for recurrent mutations to a first approximation and produces a sample-scaled mutation rate, *µT*. We treated the observed polymorphism state (i.e., presence or absence of polymorphism) at each site as a Bernoulli trial with success probability p (Fig 1C). To model the effects of sequence context, methylation, and their interaction, we assumed that the mutation rate increases linearly with methylation level, allowing both the intercept and slope to vary across the 16 4-mer sequence contexts (Fig 1B,C).

We implemented this model in a regression framework and estimated the scaled mutation rates of unmethylated and methylated CpGs in each 4-mer context using the gnomAD polymorphism data (S1 Table). In terms of scaled mutation rates, the rank order of the 16 4-mer contexts was markedly different between unmethylated and methylated states, demonstrating a strong interaction between the methyl group at the focal cytosine and flanking bases on CpG mutation rate (Fig 1D; S1B Fig). Similar analysis of polymorphisms from the 1KG project [37] yielded highly concordant context effects for methylated sites, although unmethylated sites showed slightly greater divergence (S2B Fig).

Additionally, we compared our estimates with those predicted by previous mutation models based on similar polymorphisms datasets. Carlson et al. (2018) estimated mutation rates for all unique 3-mer, 5-mer and 7-mer contexts based on extremely rare variants without considering methylation level of CpG sites. We found strong concordance between their estimates and our “methylation-weighted” estimates for each 4mer context (S3A Fig; see Methods). In turn, our estimates for unmethylated and methylated CpG sites are also in good concordance with 3-mer estimates from the gnomAD mutation models [7,36] for CpG sites in the lowest and highest methylation bins (S3B,C Fig).

To further validate our predicted mutation rates, we compared our estimates with mutation rates estimated using an independent published dataset: de novo mutations (DNM) determined by genome sequences of 5,420 parent-offspring trios [43]. Next, we identified sites with methylation level <10% and >90% in sperm as proxies for unmethylated and methylated sites, respectively. We then compared the observed DNM rates in different 4-mer contexts against our estimated scaled mutation rates for unmethylated and methylated CpGs. These comparisons showed strong correlations (Pearson’s r=0.82 and 0.97), and the two sets of estimates are approximately proportional, validating that our framework can reliably recover context-dependent *de novo* CpG mutation rates from polymorphisms.

### Independent effects of upstream and downstream bases

By organizing the 4-mer contexts by their 5’ and 3’ bases in rows and columns and visualizing inferred scaled mutation rates in heatmaps, we observed clear marginal effects of each flanking nucleotide on the mutation rates of unmethylated and methylated CpGs (Fig 2A). Specifically, we found that a 5’A has a strong, positive marginal effect on CpG mutability regardless of methylation status, which recapitulates previously reported hypermutable CpG motifs featuring an A upstream to the focal C [1,8,34,42]. We also observed a negative effect of 5’G on unmethylated CpG mutability and a strong negative effect of 5’T on methylated sites. For the downstream base, 3’C is associated with lower mutability at unmethylated CpGs, while 3’T exerts a negative impact on methylated cytosines. These inferred marginal effects of flanking bases are all replicated qualitatively using 1KG polymorphism data (S2C Fig).

**Fig 2.**
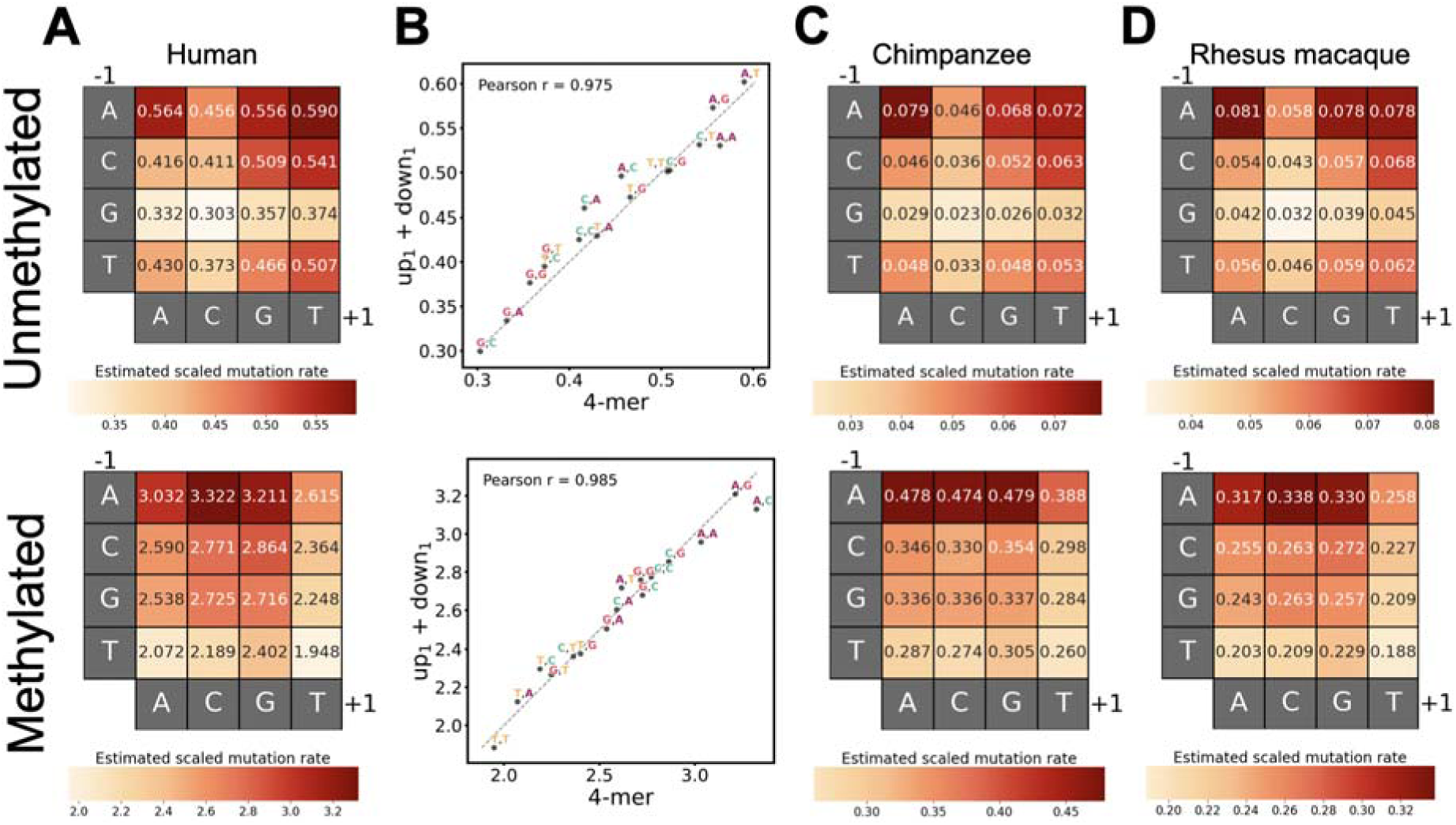
Independent effects of upstream and downstream nucleotides on CpG mutation rates and shared context effects across primates. **A.** Scaled mutation rates in humans estimated from the 4-mer model, which assumes interactions between the upstream and downstream bases. **B.** Concordance between human mutation rates estimated from the 4-mer model and from the up_1_+down_1_ model, which assumes independent effects of upstream and downstream bases. Dashed line indicates relationship. Points are labeled by the nucleotides upstream and downstream of the CpG, with error bars representing 95% confidence intervals for the estimates on both axes. **C, D.** Same as **A** but for chimpanzee and rhesus macaque. Each heatmap shows relative mutation rates for that panel; color scales are not comparable across panels.

These consistent marginal effects—where upstream bases have similar effects regardless of the downstream bases and vice versa— suggested that the flanking bases on the two sides of the CpG act largely independently. We therefore asked to what degree the mutability of each 4-mer context could be captured by the combined effect of the upstream and downstream bases. To test this, we explicitly defined a model where the upstream and downstream nucleotides independently affect the mutability of the focal cytosine (hereafter referred to as the up_1_+down_1_ model) (Table 1). This model contrasts with the 4-mer model presented above, which implicitly allows for interactions between the two flanking bases.

**Table 1.** Descriptions of the various models specified in our regression framework.

| Model | Formula | Variable | Variable description | Assumption |
| --- | --- | --- | --- | --- |
| All | | $m$ | Methylation level as measured in sperm | |
| 4-mer | $4mer * m$ | $4mer$ | 4-mer including the focal CpG and one nucleotide on either side | Each 4-mer context may have unique baseline mutation rate and methylation effect |
| $up_1+down_1$ | $U1 + D1 + m + U1:m + D1:m$ | $U1$ | Nucleotide one position upstream of focal C | Each nucleotide upstream or downstream to CpG affects the baseline mutation rate and mutagenic effect of methylation independently |
| | | $D1$ | Nucleotide one position downstream of focal G | |
| 6-mer | $6mer * methylation$ | $6mer$ | 6-mer including the focal CpG and two nucleotides on either side | Each 6-mer context may have unique baseline mutation rate and methylation effect |
| $up_{21}+down_{12}$ | $U21 + D12 + m + U21:m + D12:m$ | $U21$ | Two nucleotides upstream of focal C | Each dimer upstream and downstream to CpG may have a unique baseline mutation rate and methylation effect |
| | | $D12$ | Two nucleotides downstream of focal G | |
| $up_2+up_1+down_1+down_2$ | $U2 + U1 + D1 + D2 + m + U2:m + U1:m + D1:m + D2:m$ | $U2$ | Nucleotide two positions upstream of focal C | Each nucleotide up to two positions upstream or downstream to CpG affects the baseline mutation rate and mutagenic effect of methylation independently |
| | | $U1$ | Nucleotide one position upstream of focal C | |
| | | $D1$ | Nucleotide one position downstream of focal G | |
| | | $D2$ | Nucleotide two positions downstream of focal G | |

We found that the scaled mutation rates predicted by the up_1_+down_1_ model are highly concordant with those from the 4-mer model (Pearson’s r=0.98 for unmethylated and r=0.99 for methylated CpG; Fig 2B), confirming largely independent effects of the upstream and downstream nucleotides. Statistically, although the 4-mer model significantly outperforms the up_1_+down_1_ model based on Akaike Information Criterion (AIC) and Bayesian Informational Criterion (BIC), the gain in variance explained is small (25.18% vs 25.14%) (S2 Table). This suggests that the simpler up_1_+down_1_ model largely captures variation in CpG mutation rates, despite having fewer than half the parameters (14 vs. 32).

### Effects of sequence context on CpG mutation rate in other primate species

We then applied our regression framework to two additional primate species, chimpanzee and rhesus macaque, and observed patterns highly similar to those found in humans (Fig 2C and Fig 2D; S1 Table). First, the 4-mer context effects on mutation rate differ between methylated and unmethylated states (S4 Fig). Second, the mutation rates predicted by the up_1_+down_1_ model and 4-mer model are highly concordant, reaffirming the largely independent effects of upstream and downstream bases on focal CpG mutability (chimpanzee: Pearson’s r=0.96 and 0.99 for unmethylated and methylated CpG, respectively; rhesus macaque: Pearson’s r=0.98 for both; S4 Fig). Furthermore, most marginal effects of flanking bases are conserved across all three studied primates. In each species, a 5’A increases the mutation rate of both unmethylated and methylated CpGs, a 5’G or 3’C decreases the mutation rate of unmethylated CpGs (Fig 2A,C,D), and a 5’T or 3’T decreases the mutation rate of methylated CpGs. Together, these findings suggest strong conservation in context-dependent mutation patterns of CpG sites across these primate species.

Despite these broad similarities, clear inter-species differences emerge in the context-dependency of CpG mutation rates. Due to greater estimation uncertainty, inter-species correlations of point estimates are difficult to interpret for unmethylated sites. However, for methylated sites where estimation resolution is high, similarity in context effects is surprisingly highest between human and rhesus macaque, while either species shows lower concordance with chimpanzee (Fig 3). Specifically, the greatest inter-species divergence occurs between humans and chimpanzees (Pearson’s r = 0.914), with the most pronounced discrepancies observed in 4mer contexts with a downstream cytosine: a 3’C has a strong positive marginal effect in humans and rhesus macaques but a relatively lower effect in chimpanzees (Fig 2A, C, D; Fig 3A). Together, these results suggest that context-specific CpG mutability, particularly at methylated sites, may have shifted during primate evolution (Fig 3B,C).

**Fig 3.**
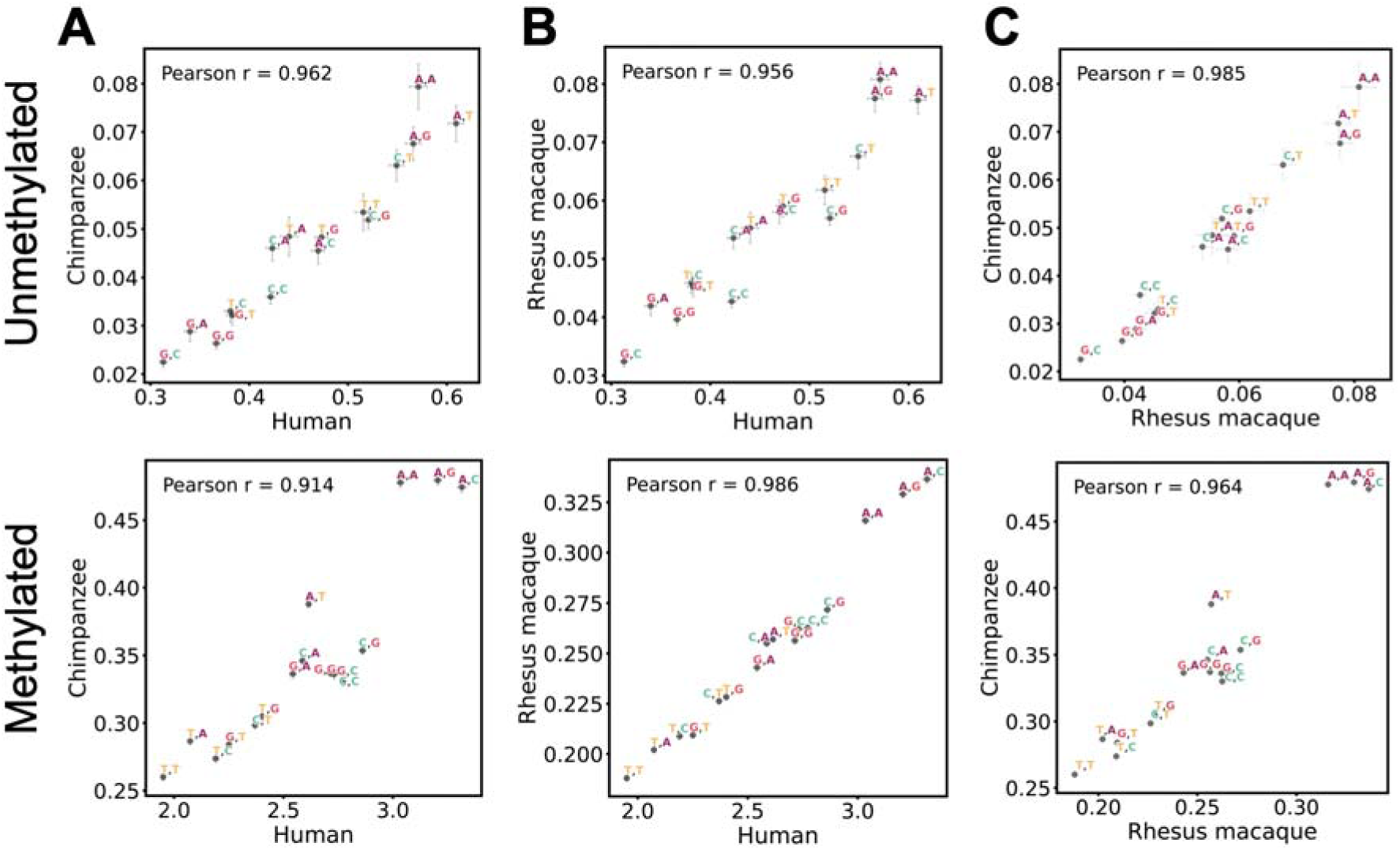
Similarity and differences in context effects on CpG mutability across primates. **A.** Comparison of scaled mutation rate estimates between human and chimpanzee at unmethylated and methylated sites, with error bars representing 95% confidence intervals for the estimates on both axes. **B.** Same as **A.** for comparisons between human and rhesus macaque. **C.** Same as **A.** for comparison between rhesus macaque and chimpanzee.

### Mutational asymmetry within the same CpG dinucleotide

Given that the upstream and downstream nucleotides have asymmetric influences on the mutability of the focal cytosine, we wondered whether the two adjacent C:G base pairs within the same CpG dinucleotide differ in mutation rate. To test this, we compared mutation rates of reverse complement 4-mer contexts (e.g., ACGC and GCGT). Since our modeling and analysis always measure mutation rate at the cytosine position of each CpG, each pair of reverse complement contexts correspond to the two adjacent C:G base pairs within the same CpG dinucleotide (Fig 4).

**Fig 4.**
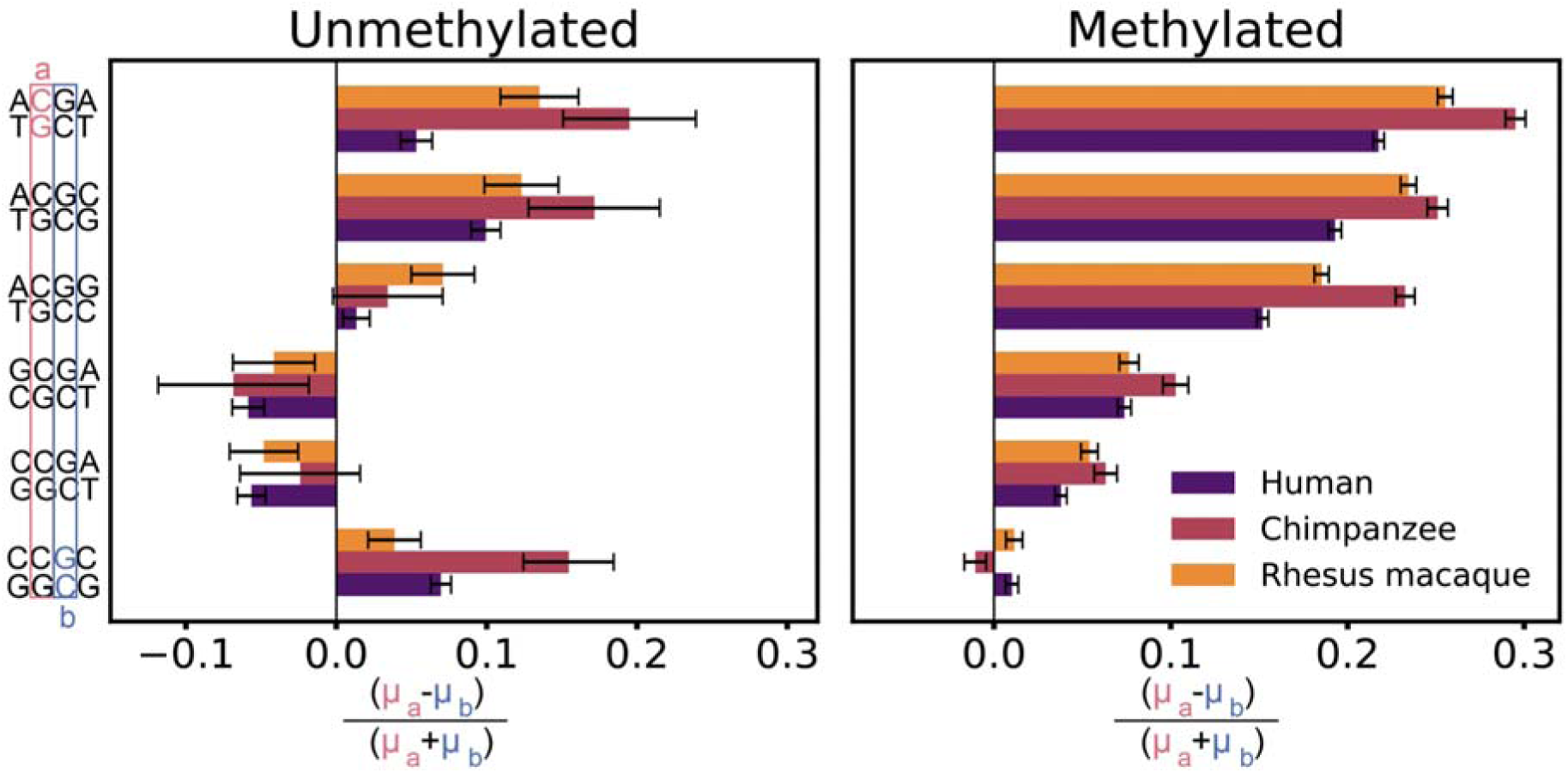
Mutational asymmetry within a CpG site. Each bar represents a pair of reverse complement 4-mer contexts (e.g., A**C**GC and G**C**GT) with a different focal cytosine for mutation rate estimation, with color indicating the species (human, chimpanzee, or rhesus macaque). Within the same CpG dinucleotide, adjacent C:G base pairs *a* and *b* often show different mutation rates in both unmethylated and methylated states. Error bars represent 95% confidence intervals calculated using the context-specific scaled mutation rate estimates from our model (S1 Table) and assuming propagation of uncertainty.

In humans, we found significant mutational asymmetry between the two adjacent base pairs at both methylated and unmethylated sites across most context pairs (Fig 4). The direction of this asymmetry is concordant for unmethylated and methylated CpGs in four out of six 4-mer context pairs. Notably, a cytosine directly flanked by an upstream A is consistently more mutable than the neighboring G:C base pair within the same CpG dinucleotide, reflecting the strong mutagenic effect of 5’A (Fig 2). For the CCGC:GCGG pair, we observed significant asymmetry for unmethylated CpGs, with the C:G base pair flanked by an upstream C:G being more mutable; the asymmetry is in the same direction but much weaker for methylated CpGs. For the CCGA:TCGG and GCGA:TCGC pairs, methylation status determines the direction of mutational asymmetry: the G:C base pair directly flanked by a downstream A:T is more mutable than its counterpart when unmethylated but less mutable when methylated.

Consistent with the findings in humans, adjacent C:G base pairs within the same CpG also differed in mutability in chimpanzees and rhesus macaques for most contexts (Fig 4). When significant asymmetry is observed for a context pair, the direction of asymmetry is usually consistent across all three species at both unmethylated and methylated sites, reflecting the well-conserved effects of certain flanking bases (Fig 2). The only exception occurs at methylated sites in CCGC:GCGG, where chimpanzee shows asymmetry in the opposite direction than human and rhesus macaque, although the magnitudes of asymmetry are weak in all three species. This flip in asymmetry echoes the reduced marginal effect of 3’C in chimpanzee (Fig 2C).

### Effects of expanded sequence context on CpG mutation rate in primates

Beyond the immediate flanking bases, the expanded sequence context is a known predictor of mutation rate across the genome [8,33,34]. We leveraged our regression framework to analyze 6-mer contexts, incorporating nucleotides two positions upstream and downstream. The 6-mer model implicitly assumes all possible interactions between the four flanking nucleotides. Consistent with our observations for 4-mer contexts, effect of the same 6-mer context differs between unmethylated and methylated sites, confirming interactions between flanking nucleotides and the methyl group (Fig 5).

**Fig 5.**
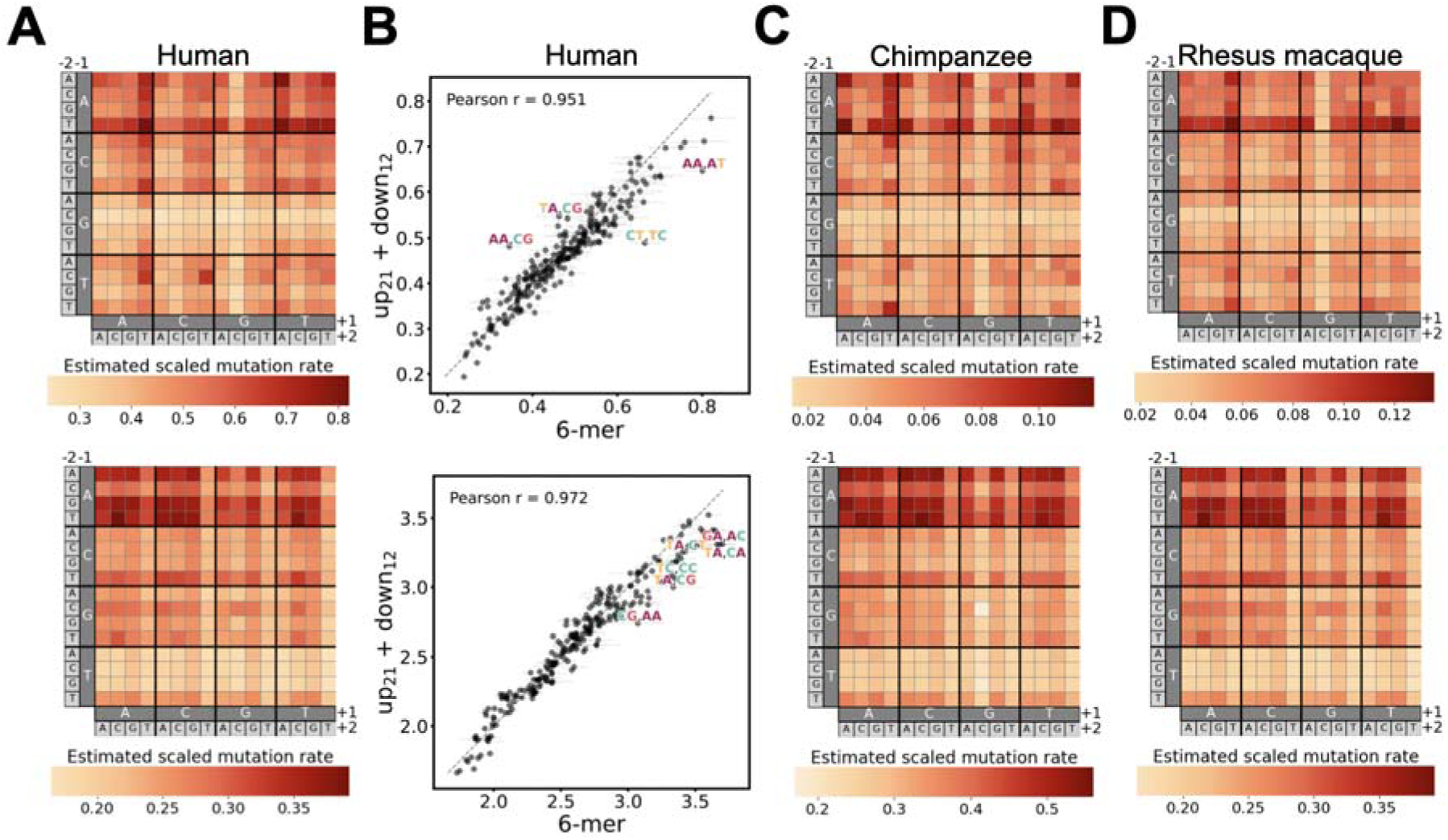
Shared and diverged effects of upstream and downstream sequences on CpG mutability across primates at 6-mer resolution. **A.** Scaled mutation rates estimated from the 6-mer model for human at unmethylated (upper) and methylated sites (lower). **B.** Concordance between scaled mutation rates estimated from the 6-mer model and the up_21_+down_12_ model, which assumes interactions only between bases within the upstream and downstream dimers. Dashed line indicates relationship. Points are labeled by the dimers upstream and downstream of the CpG. Only contexts for which the difference between the two model estimates exceeds 2.5 standard deviations from the mean difference are labeled. Error bars represent 95% confidence intervals for the estimates. **C, D.** Same as **A** but for chimpanzee and rhesus macaque. Each heatmap shows relative mutation rates for that panel; color scales are not comparable across panels.

Exploring the marginal effects of upstream and downstream dimers in humans revealed more refined context-dependencies (Fig 5A). Echoing the strong mutagenic effect of upstream A detected at the 4-mer scale, an upstream TA dimer has the strongest, positive marginal effect at unmethylated sites, whereas an upstream GA or TA dimer is mutagenic for methylated CpGs. Notably, at methylated sites, a T at the downstream +2 position (T_+2_) is consistently associated with low mutation rates, regardless of the intermediate +1 nucleotide.

The row- and column-wise patterns in the 6-mer heatmaps suggested that the upstream and downstream dimers may act independently. We tested this by comparing the full 6-mer model against a simpler “up_21_+down_12_” model, which only includes interactions between adjacent bases within each upstream or downstream dimer but no interaction between upstream and downstream sequences. We again found high concordance in inferred scaled mutation rates predicted by the two models (Pearson’s r=0.95 and 0.97 for unmethylated and methylated CpG, respectively; Fig 5B). Although the 6-mer model significantly outperforms the up_21_+down_12_ (S2 Table), the additional variance explained is small (25.52% vs 25.43%).

We then tested an even simpler model (up_2_+up_1_+down_1_+down_2_), where all flanking nucleotides contributed independently to CpG mutability. This further simplified model showed weaker concordance with the up_21_+down_12_ model (Pearson’s r=0.93 and 0.94 for unmethylated and methylated CpG, respectively; S5 Fig), suggesting that interactions within the upstream or downstream dimers contribute more to CpG mutation rate variation. Although more complex models statistically outperform simpler ones (S2 Table), they involve many more parameters that require large datasets to estimate reliably. Interestingly, the up_2_+up_1_+down_1_+down_2_ model, despite having fewer parameters and assuming no interactions at all, outperforms the 4mer model that considers shorter contexts but full interaction among base pairs (variance explained 25.43% vs 25.18%; S2 Table), pointing to the significant influence of extended sequence on mutation rates [8,33,34]. Overall, the up_21_+down_12_ model seems to offer the best trade-off by capturing the bulk of context effects with an order of magnitude fewer parameters than the 6-mer model (62 vs. 512).

Extending this analysis to include chimpanzee and rhesus macaque identified many conserved and some species-specific 6-mer effects. We found that the upstream CG dimer and downstream GC dimer are consistently associated with low mutation rates at unmethylated sites; a 5’TA dimer has a strong, positive effect on both methylated and unmethylated CpG mutability in all three species (Fig 5A,C,D). However, at methylated sites, differences in context effects become more noticeable among the three species. Notably, the striking negative effect of downstream T_+2_ on methylated CpG in humans is seen in rhesus macaques but slightly edged by a strong negative effect of the downstream GC dimer in chimpanzees. Across all three species, the most complex 6-mer model always performs considerably better than simpler models, with variance explained increasing with model complexity (S2 Table).

### Effects of sequence context on CpG mutation rate in an insect species with minimal DNA methylation

Our analysis of primate polymorphisms treated methylation as a continuous variable, but sites classified as “unmethylated” based on sperm bisulfite sequencing may still experience methylation in other tissues or developmental stages. To approximate a truly unmethylated state, we extended our analysis to the silkworm (*Bombyx mori*), a species with less than 1% methylation genome-wide, largely in gene bodies [44]. We modified our regression framework to isolate sequence context effects by eliminating the methylation component entirely (S1 Table). As in our primate analyses, we also tested simpler additive models without higher-order interactions between flanking bases.

In *B. mori*, mutation rate estimates from the 4-*mer* model and the up_1_+down_1_ model were highly concordant with nearly identical AIC (Pearson’s r = 0.98; S6A Fig; S2 Table), again indicating that flanking bases influence CpG mutability largely independently. Crucially, some of the marginal effects of flanking nucleotides in the silkworm mirror those found in primates at unmethylated sites: a 5’A increases while a 3’C decreases CpG mutability (S6B Fig; Fig 2). These shared patterns support deep conservation in context effects on CpG mutability that are independent of DNA methylation.

## Discussion

Here, we present a regression framework that enables estimation of CpG mutation rates in different 4-mer and 6-mer contexts at unmethylated and methylated states, separately. Our model introduces two key innovations: treating methylation level as a continuous predictor and explicitly modeling recurrent mutations. This approach avoids the arbitrary binning of methylation levels into discrete categories (e.g., high, medium, low) and reduces systematic underestimation of mutation rates at hypermutable sites.

While sperm bisulfite sequencing provides a practical approximation of methylation levels in the male germline, it has clear limitations. Sperm-based methylation data may not accurately reflect levels in spermatogonial stem cells, and they fail to capture methylation profiles in other cell types relevant to germline mutagenesis (e.g., female germ cells, early embryonic tissues). A particular concern is that some sites classified as “unmethylated” in our analysis may in fact be methylated in these other contexts. To strengthen our findings on unmethylated sites, we applied our model to silkworm, a species with very low levels of DNA methylation that is largely concentrated in gene bodies [44]. The recovery of consistent effects of certain flanking bases (e.g., positive effect of upstream A) in this system supports that these mutation patterns likely reflect intrinsic properties of unmethylated CpG in different DNA sequences rather than artifacts of transient methylation and/or extrapolation.

Unlike several prior mutation models, we explicitly account for recurrent mutations on observed polymorphisms. At hypermutable sites in large samples, recurrent mutations make the observed polymorphism rate a poor linear proxy for mutation rate (Fig 1A). To address this, we used a standard exponential transformation to relate polymorphism probability to mutation rate, assuming a constant total length of the coalescent tree (T) across the genome. However, genealogical histories certainly vary among genomic loci due to recombination and coalescent stochasticity, as well as systematic factors like background selection, all of which cause heterogeneity in T [45,46]. This assumption is especially problematic for species with very small sample sizes, such as chimpanzees (S3 Table), where sampling stochasticity in the coalescent process is greater. The over-simplifying assumption of constant T may distort our quantitative estimates of context-specific mutation rates and introduce biases in between-species comparisons. Nonetheless, these factors likely average out across the hundreds of thousands of CpG sites within each context, so the heterogeneity in *T* is unlikely to alter the relative ranking of contexts in the presence of substantial differences in mutability. We thus focus on qualitative rather than quantitative comparisons across contexts and species. We also note that residual variance in mutation rate that arise from mis-specification or unmodeled biological determinants in our modeling framework could introduce uncertainty in the inferred ranking of sequence contexts, especially those with similar mutation rates.

Despite limitations, our analysis clearly demonstrates that CpG mutation rate variation is shaped by the interaction between the local sequence context and methylation at the focal cytosine. Specifically, the effects of flanking sequences are distinct on unmethylated and methylated cytosines but conserved across species, supporting the view that these two cytosine states act as fundamentally different mutational substrates. Our analysis also demonstrates that the upstream and downstream flanking sequences exert largely independent influences on mutability, a pattern that holds for both 4-mer and 6-mer sequence contexts. Lastly, we identify several strong marginal effects of adjacent nucleotides on CpG sites, including a strong mutagenic effect of an upstream A on both unmethylated and methylated CpGs, negative effects of 5’CG and 3’GC dimers on unmethylated CpGs, and a positive effect of 5’TA dimer on unmethylated CpGs.

Importantly, the context and methylation effects we report here can help generate hypotheses and guide interpretations about the underlying mechanisms of germline CpG>TpG mutagenesis. For example, recent work has shown that transcription factor (TF) binding sites may act as mutation hotspots by interfering with DNA repair [32], with TF motifs typically spanning 6-10 nucleotides [47]. By contrast, our results indicate that CpG mutability is primarily explained by largely independent effects of upstream and downstream sequences. This suggests that the molecular mechanisms underlying context effects may operate in a modular fashion—upstream and downstream sequences likely influence distinct steps in mutagenesis rather than requiring complex long-range structural interactions. This argues against TF occupancy as a dominant driver of context-dependent CpG mutation rate variation in the germline. Future work should test whether this modularity extends to CpG transversions or other single-nucleotide substitutions. If confirmed, this modularity has practical implications: simpler additive or multiplicative models can achieve nearly the same predictive power as fully parameterized models while remaining interpretable and computationally tractable.

Notably, an upstream A substantially increases mutability at both unmethylated and methylated sites relative to other bases, which recapitulates previously reported hypermutable motifs [1,8,34] and appears conserved across species, including chimpanzee, rhesus macaque, and even silkworm. This deep conservation suggests that the ACG motif has intrinsic biophysical properties—perhaps related to DNA shape—that elevate CpG mutation rates independent of methylation status. Intriguingly, previous work shows that C>T mutation rates are also elevated at non-CpG sites flanked by an upstream A [34], indicating that hypermutability may be a general property of ACN sequence, regardless of CpG or non-CpG sites, which calls for investigation into the mechanism.

In contrast, an upstream G does not elevate mutation rate at methylated sites and consistently reduces mutation rate at unmethylated CpGs in all three primate species (Fig 2). This observation contrasts with recent reports of elevated error rate of wild-type DNA polymerase _ε_ in GCG contexts [12]. Together, these findings suggest that although deamination-independent replication-related CpG>TpG mutations occur, they cannot explain much of the context-dependent mutation rate variation and are unlikely to represent the major source of germline CpG mutations.

Cross-species comparisons of CpG mutation patterns offer additional mechanistic insights, revealing that divergence in context effects does not strictly follow known phylogenetic relationships. While correlations between point estimates at unmethylated sites may be difficult to interpret due to estimation uncertainty, a clearly surprising pattern emerges at methylated sites, where our model should have high resolution: humans and rhesus macaques exhibit remarkably high similarity (Pearson’s r = 0.986), while either species shows lower concordance with the chimpanzee (Pearson’s r = 0.914 and 0.964, respectively). At the current stage, we cannot fully rule out technical factors related to the chimpanzee polymorphism data, making it essential to replicate these findings using independent chimpanzee datasets or alternative methodologies. If this divergence is indeed robust, the most parsimonious explanation is rapid evolutionary shifts in the context-dependency of 5mC mutation rates along the chimpanzee lineage. Such shifts may reflect changes in the context-specificity of enzymes involved in active demethylation, such as the TET family [43], or in the repair of deamination products by proteins like thymine DNA glycosylases (TDGs) [44].

## Methods

### Extracting CpG sites

We retrieved the hard-masked reference genomes for human (hg38), chimpanzee (panTro5), rhesus macaque (Mmul10), and silkworm [48] from UCSC genome browser and Silkbase (see Data sources). To minimize the effects of selection, we excluded genic regions based on NCBI RefSeq annotations for all species studied; for human, we also removed phylogenetically conserved regions defined by *phastcons* (see Data sources). For the primate species, we also obtained the inferred reconstructed ancestral genome from the 10-primate EPO alignment [49] mapped onto the human (hg38), chimpanzee (panTro5), and rhesus macaque (Mmul10) genome. We then intersected the hard-masked reference genome for each species with the corresponding ancestral reference and did all downstream analysis on hard-masked sites that were also present in the ancestral genome.

Then, we scanned the genome to identify every occurrence of “CG” and recorded the position of the dimer along with the flanking sequences. Each CpG dinucleotide was treated as two independent cytosines: one on the forward strand in the extracted context and the other on the reverse strand in the reverse complement sequence context. Based on flanking sequences, we extracted 4-mer and 6-mer contexts as needed. To test whether any sites lost after intersection could bias our estimates, we compared results based on all sites in hard-masked reference genomes to those in intersected regions and observed largely concordant context-specific mutation rate estimates (S7B Fig, S4 Table). The intersected genome yielded improved model performance, as measured by the proportion of variance explained. Accordingly, all analyses and results in the main text are based on the intersected regions.

### Data sources

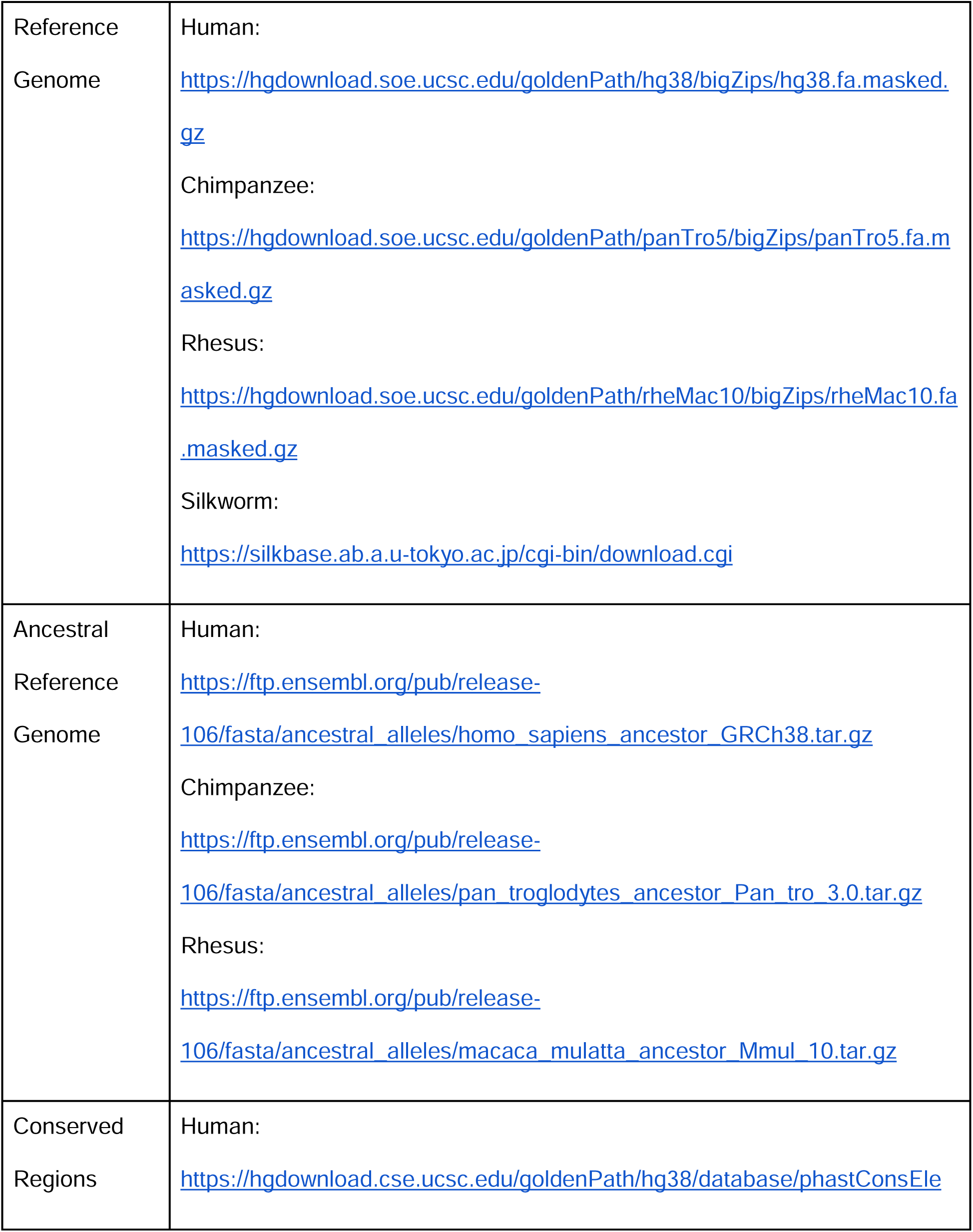
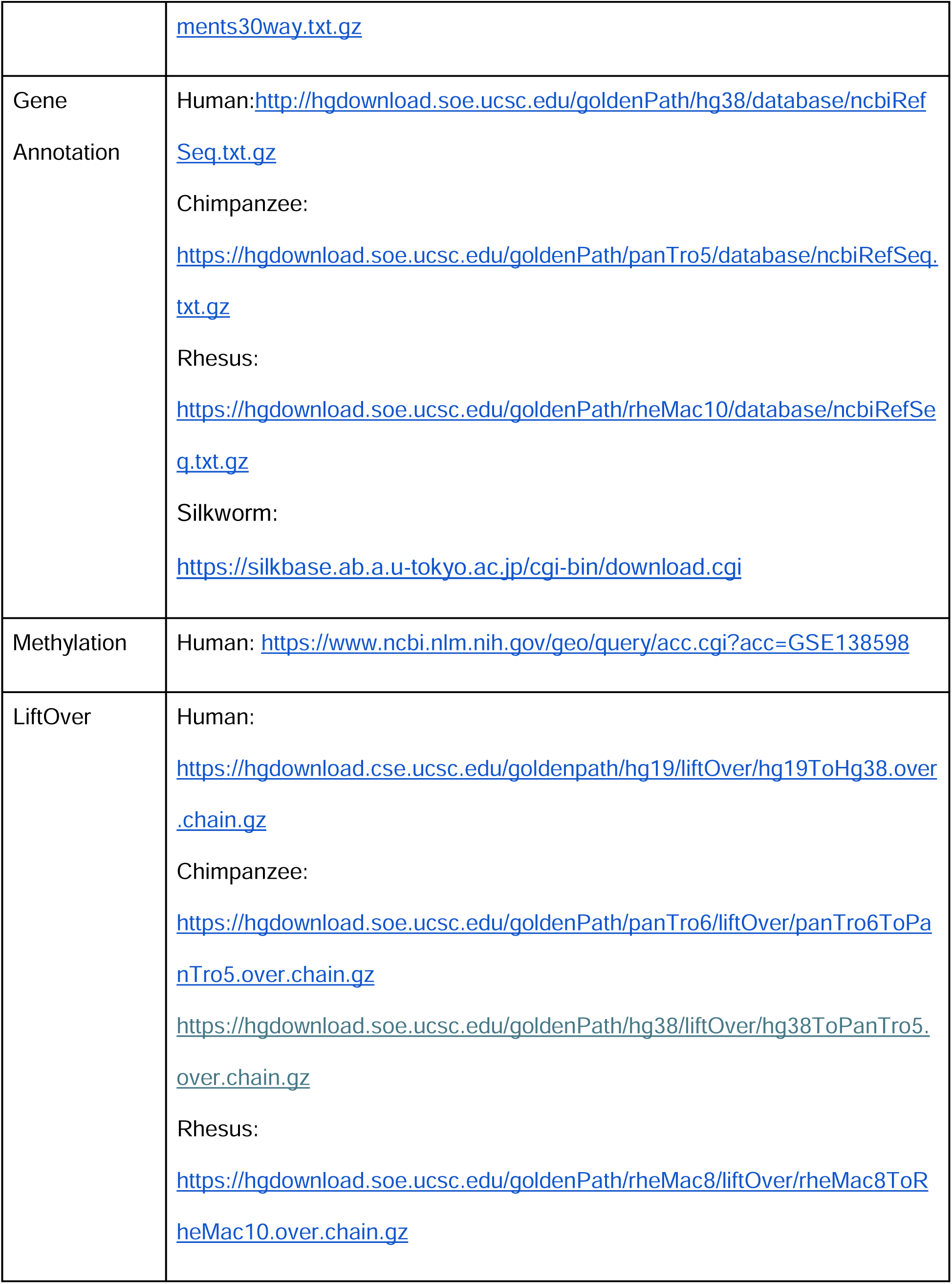
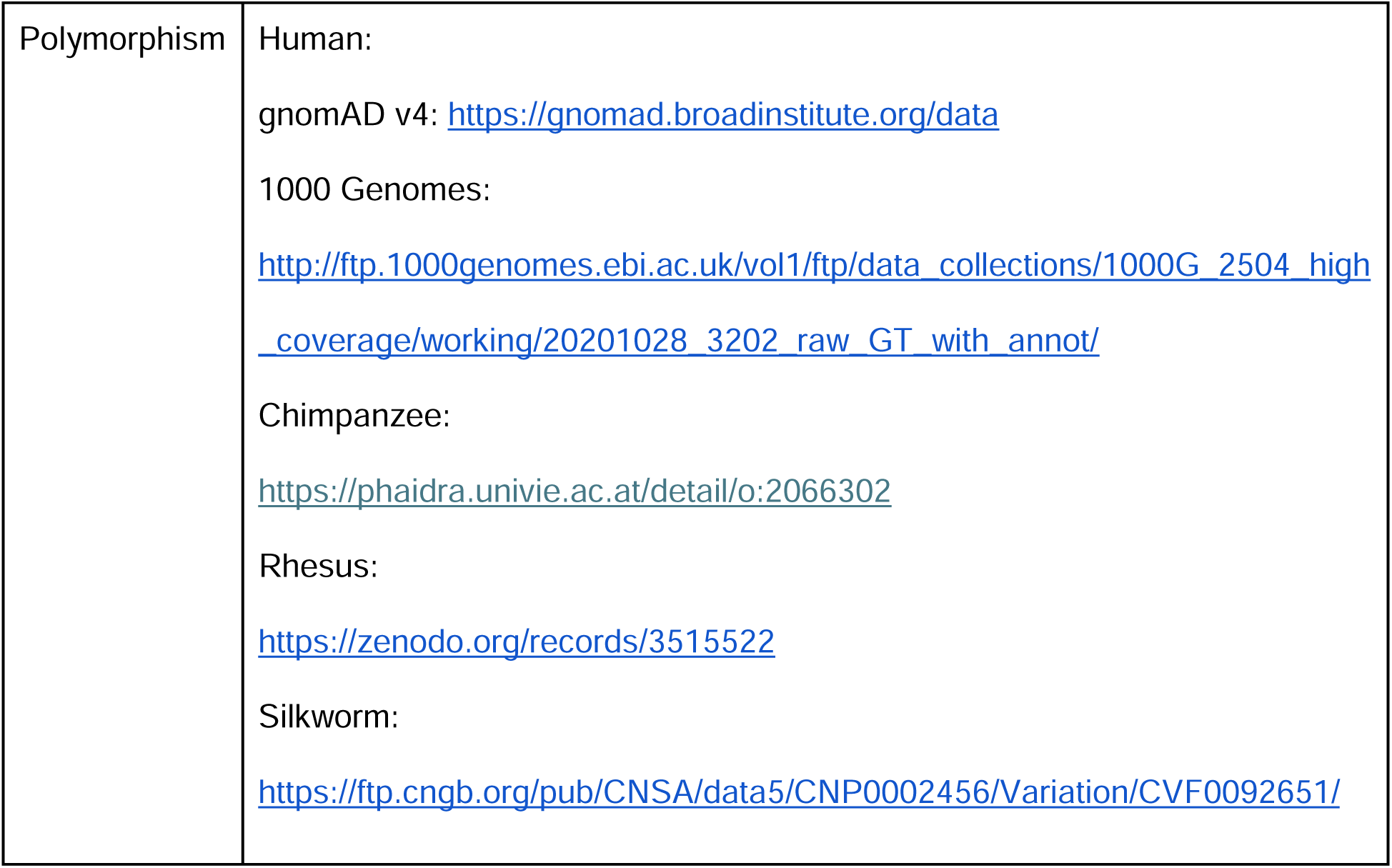

### Methylation data

For human, we downloaded bisulfite sequencing data from Chen et al. [38], which reported sperm WGBS data from nine healthy individuals. When a C/T SNP exists at a CpG site, methylation estimates are biased in individuals carrying one or two T alleles, as the T allele will be mistaken for a bisulfite-converted unmethylated C. For instance, even if the CpG allele is “fully” methylated in sperm, the presence of the TpG allele results in incorrect reporting of only 50% methylation in heterozygotes and 0% methylation in T/T homozygotes. To reduce this bias, we calculated the average methylation level at each CpG by pooling methylated and unmethylated read counts across samples. This approach effectively reduces biases for most CpGs where the derived T allele is rare in the nine individuals but still underestimates methylation levels at sites where the T allele is at high frequency. Additionally, this pooling strategy assumes that methylation patterns are relatively consistent across individuals at most CpG sites, which may not hold for regions with high inter-individual variability or imprinted loci. We then lifted over the average methylation level from hg19 to hg38.

To test whether hydroxy-methylation (5hmC), which cannot be distinguished from 5mC by traditional bisulfite sequencing, distorts our estimation of mutation rates for unmethylated cytosine and 5mC, we used published 5hmC calls from TET-assisted bisulfite sequencing in human embryonic stem cells [50]. In this dataset, about 2.1% of CpGs in putatively neutral regions of the human genome have a recorded non-zero 5hmC level, with an average level of 21%. We re-ran our model after excluding all sites with any non-zero 5hmC signal, and found that the context-dependent mutation rate estimates were nearly perfectly concordant with those estimated using all sites (S8 Fig), suggesting that hydroxy-methylated sites mis-labeled as methylated sites do not have an appreciable effect on the estimation of context-specific mutation rates at unmethylated cytosine and 5mC.

For chimpanzee and rhesus macaque, we retrieved testes EM-Seq methylation calls averaged between two individuals and corrected for genotype (A. Stolyarova, personal communications) at each CpG dinucleotide. Chimpanzee methylation data was lifted over from panTro6 to panTro5.

### Filtering and polarization of polymorphism

We downloaded autosomal polymorphism data for human [36,37], rhesus macaque [51], chimpanzee [52], and silkworm [53] (see Data sources). For rhesus macaque, variant coordinates were lifted over from Mmul8 to Mmul10. For chimpanzee, variant coordinates were lifted over from hg38 to panTro5. Singletons were excluded in the silkworm dataset to reduce false-positive variant calls [54]. For human, we filtered singletons to positions with single-read Umap100 mappability equal to 1, ensuring that variants were supported by uniquely mappable 100-bp reads and minimizing spurious rare variants due to alignment errors [55].

We polarized SNPs based on allele frequency, defining the major allele as the ancestral state and the minor allele as the derived state (hereby referred to as MAF polarization). For silkworm, we used *est-sfs* [56], which estimates the probability that the major allele of the focal species (*Bombyx mori*) is the ancestral state using polymorphism data from the focal species and alignment with a closely related outgroup (*Bombyx mandarina*). We annotated each SNP with the type of substitution and retained only C>T and G>A mutations at CpG sites (extracted as described above, see S3 Table).

We also used an alternate polarization method for the primate species, by inferring the ancestral allele based on the 10 primate EPO ancestral genome reconstruction. While we found similar context-specific mutation rate estimates using this method (S7A Fig), the MAF polarization method generally yielded better model performance in terms of proportion variance explained across all primate species, regardless of the CpG annotation method used (S4 Table). Therefore, we decided to use MAF polarization in our main analysis.

The number of genotyped samples in a polymorphism dataset can vary across sites and can systematically differ by context thus confounding the estimation of context-specific rates. To address this, we examined the distribution in sample coverage across contexts in the polymorphism datasets for the three primate species and found no significant differences (S9 Fig).

In addition, GC-biased gene conversion (gBGC) can have a profound impact on the frequency of C>T polymorphisms [57]. Since the strength of gBGC depends on population sizes and the specific effects are also sample-specific, differences in effective population sizes and sample sizes across species can bias the scaled mutation rates. However, the strength of gBGC is not expected to differ substantially across contexts and thus unlikely to lead to significant biases in the rankings of contexts. To address this directly with data, we stratified CpG sites by recombination rate [58] into four bins covering roughly the same number of CpG sites: high (> 0.389), mid1 (≤ 0.389), mid2 (≤ 0.026), and low (≤ 4.877e-05). We then estimated the mutation rate for each bin separately. Consistent with our expectation, we find strong correlations between estimates from all sites and those of each recombination bin for methylated CpGs (Pearson’s r > 0.99) and slightly lower correlation for unmethylated CpGs (Pearson’s r>0.97). The reduction of correlation at unmethylated sites may be explained by noisier point estimates due to smaller numbers of sites with low methylation levels (S10 Fig).

### Modeling effects of methylation and sequence contexts on CpG mutation rate

To quantify the effects of local sequence context and methylation level on CpG mutability, we modeled the mutation rate *µ_i_* at a given cytosine i (within a CpG dinucleotide) as a linear function of its methylation level *m_i_*, allowing both the intercept and slope to vary with sequence context:

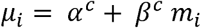

where, *α^c^* represents the baseline mutation rate for context *c* and *β3^c^* denotes the effect of full methylation within that context.

For a site with mutation rate of *µ* per generation, the probability of having polymorphism in a sample was modeled as *p* = 1 - *e*^-*µT*^, where T is the total coalescent branch length (in generations) connecting all sampled genomes. For simplicity, we assumed that *T* was constant across all CpG sites in non-genic, non-conserved regions. The presence or absence of polymorphism at each site was then modeled as a Bernoulli trial. i.e., drawn from a binomial distribution *Binom*(1, *p*) with *p* = 1 - *e*^-*µT*^.

### Regression framework

To implement this model, we used the *glm* function in R, specifying the binomial family and a custom link function, which maps the polymorphism probability (*p*) to a linear predictor *η* = -*µT* = log (1 - *p*) as follows:

- Link function: *link(p)* = log (1 - *p*)
- Inverse link function: link^-l^(*η*) =1 - *e^η^*
- Derivative of the inverse link: 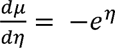

We fitted the generalized linear model to site-level data, where each record corresponded to one base pair in a CpG site annotated with its methylation level, sequence context, and a binary indicator of polymorphism status (0 = monomorphic, 1 = polymorphic). The regression model was specified differently depending on the context length and type of interaction, as described in Table 1.

After fitting the regression models to the polymorphism data, we estimated the scaled mutation rates (µT) for each sequence context and methylation status (*x*=1 for methylated; *x*=0 for unmethylated) using the predict() function in R with type = “link” to obtain the linear predictor (η).

To evaluate model performance, we extracted summary statistics from each fitted regression and compared models using criteria that balance between model fit and complexity, namely the Akaike Information Criterion (AIC) and the Bayesian Information Criterion (BIC), which is preferred for larger datasets such as ours. We also measured explanatory power as the proportion of variance in polymorphism status explained, calculated as the difference between null and residual deviance, divided by the null deviance (S2 Table).

## Acknowledgements

We thank Anastasia Stolyarova and Molly Przeworski for kindly sharing sperm methylation data for chimpanzee and rhesus macaque before publication; Iain Mathieson, Fabian Ramos-Almodovar, and other members of the Gao and Mathieson laboratories for helpful discussions.

## Funder information

This work is supported by a Research Fellowship (**FG-2021-15702**) from the Alfred P. Sloan Foundation (https://sloan.org/) and a grant (**R35GM146810**) from the National Institute of General Medical Sciences to ZG. The funders had no role in study design, data collection and analysis, decision to publish, or preparation of the manuscript.

## Supporting Information

**S1 Fig. Interaction between the methyl group at the focal cytosine and flanking bases on CpG mutation rate. A.** Polymorphism rate for each 4-mer context, stratified by methylation levels. CpG sites were partitioned into 20 equal-width bins (ranging from [0, 0.05) to [0.95, 1]). Each point represents the proportion of SNPs among all CpG sites within a methylation bin, along with 95% confidence intervals assuming binomial sampling. Each context is represented by a color denoting the nucleotide at the 5’ position and a shape denoting the nucleotide at the 3’ position. **B.** Scaled mutation rates for each 4-mer context as estimated by the model for unmethylated and methylated sites. Points are labeled by the nucleotides upstream and downstream of the CpG, with error bars representing 95% confidence intervals for the estimates.

**S2 Fig. Concordance between mutation rates for 4-mer CpG contexts estimated from gnomAD (v4) and 1000 Genomes (1KG) polymorphism datasets. A.** Scaled mutation rate estimates for unmethylated and methylated sites for each 4-mer context using 1KG data. The contexts are sorted in descending order of mutability when unmethylated and error bars represent 95% confidence intervals for the estimates. **B.** Comparison of mutation rates estimated using polymorphisms from gnomAD and 1KG at unmethylated and methylated sites, respectively. **C.** Heatmap of scaled mutation rate estimates using the 4-mer model at unmethylated and methylated sites, using polymorphism data from 1KG. Each heatmap shows relative mutation rates for that panel; color scales are not comparable between panels. **D.** Concordance between mutation rates estimated from the 4-mer model and from the up_1_+down_1_ model using 1KG data. Points are labeled by the nucleotides upstream and downstream of the CpG, with error bars representing 95% confidence intervals for the estimates in panels B and D. The dashed line indicates x = y.

**S3 Fig. Context-specific scaled mutation rates are consistent with estimates from other published context-dependent mutation models. A.** Comparison with rates published in Carlson et al. (2018). To compare Carlson et al.’s estimates for 5-mers and 7-mers with our 4-mer and 6-mer results, we derived 4-mer and 6-mer rates as weighted averages of the mutation rate estimates for 5-mer (top) and 7-mer (bottom) contexts reported by Carlson et al., using the average methylation level of each 5-mer or 7-mer context as weights. For the expanded context comparison, a linear fit constrained through the origin is shown (dashed line). Only contexts with residuals exceeding 2.5 standard deviations from the fitted line are labeled. **B, C.** Comparison with rates published from gnomAD mutation rate models (Karczewski et al., 2020; Chen at al., 2024) for unmethylated (B) and methylated CpGs (C). To facilitate comparison, we computed the mutation rate for each 3-mer as the mean of estimates of the four corresponding 4-mer contexts sharing the same upstream base (e.g., rate for ACG is the mean of the rates of ACGA, ACGC, ACGG and ACGT). In all panels, points are labeled by the nucleotides upstream and downstream of the CpG, with horizontal error bars representing 95% confidence intervals for the model estimates.

**S4 Fig. Independent effects of upstream and downstream bases on CpG mutability in chimpanzee and rhesus macaque. A.** Concordance between mutation rates as estimated from 4-mer model and from up_1_ + down_1_ model which assumes independent effects of upstream and downstream bases in chimpanzee for unmethylated and methylated sites. **B.** Same as (A), for rhesus macaque. Points are labeled by the upstream and downstream nucleotide flanking the CpG, and error bars denote 95% confidence intervals. The dashed line indicates x = y.

**S5 Fig. Concordance in mutation rate estimates between models with and without interactions within upstream and downstream dimers.** Comparison of human mutation rates as estimated from the up_21_+down_12_ model, which assumes interactions between bases within the dimer upstream and downstream to the CpG, and the up_2_+up_1_+down_1_+down_2_ model, which assumes independent effects of every flanking base. Only contexts for which the difference between the two models’ estimates is more than 2.5 standard deviations away from the average difference across all contexts are labelled. Error bars represent 95% confidence intervals for the estimates on both axes. The dashed line indicates x = y.

**S6 Fig. Independent effects of upstream and downstream bases on CpG mutability in the insect species, *Bombyx mori* (silkworm). A.** Concordance between mutation rates as estimated from 4-mer model and from up_1_+down_1_ model. Mutation rate is modelled as a function of sequence context alone. Points are labeled by the upstream and downstream nucleotide flanking the CpG, and error bars denote 95% confidence intervals. The dashed line indicates x = y. **B.** Heatmap of estimated scaled mutation rates using the 4-mer model.

**S7 Fig. Context-specific scaled mutation rate estimates are highly concordant regardless of CpG filtering or SNP polarization method. A.** Estimates obtained by polarizing SNPs using minor allele frequency (MAF) are compared to those based on the inferred ancestral allele (ANC) for CpG sites present in both the hard-masked reference and inferred ancestral genome. **B.** Estimates derived from the intersected reference genome—restricted to CpGs present in both the hard-masked reference and inferred ancestral genome—are compared to those from the masked reference genome without intersection (“Masked”); SNPs are polarized by MAF. Points are labeled by the upstream and downstream nucleotide context flanking the CpG, and error bars denote 95% confidence intervals around the estimates.

**S8 Fig**. **Scaled mutation rate estimates are nearly perfectly concordant after excluding hydroxy-methylated (5hmC) sites.** Comparison of the model’s scaled mutation rate estimates for unmethylated and methylated sites when using all sites and when excluding sites with a non-zero 5hmC level as measured in human ES-cells. Points are labeled by the upstream and downstream nucleotide context flanking the CpG, and error bars denote 95% confidence intervals around the estimates.

**S9 Fig. Callable allele coverage across 4-mer contexts in primate polymorphism datasets.** Distribution of the number of callable alleles by 4-mer sequence context in the human, chimpanzee, and rhesus macaque polymorphism dataset. Boxes show the median and interquartile range, with whiskers extending to 1.5x the interquartile range.

**S10 Fig. Context-specific scaled mutation rate estimates are consistent across recombination rate strata. A.** Comparison of scaled mutation rate estimates across all sites versus sites stratified by recombination rate, shown separately for unmethylated (top) and methylated (bottom) CpGs. Panels correspond to recombination rate quartiles: **A.** highest (“High”, > 0.389), **B** upper-middle (“Mid1”, ≤ 0.389), **C.** lower-middle (“Mid2”, ≤ 0.026), and **D.** lowest (“Low”, ≤ 4.877 × 10^-5^). The dashed line indicates x = y. Points are labeled by the upstream and downstream nucleotide flanking the CpG, and error bars denote 95% confidence intervals.

**S1 Table. Scaled mutation rate estimates for each 4-mer and 6-mer CpG context at unmethylated and methylated sites (provided as excel file).** This table contains separate worksheets for each species and model (e.g., 4-mer, 6-mer), with each sheet reporting the sequence context, methylation status, dataset-specific scaled mutation rate estimate (µT), and its associated standard error. For silkworm, we report estimates from 4-mer and up_1_+down_1_ models only, and rates are modeled as a function of context alone.

**S2 Table. Model comparison for human and non-human species.** Summary statistics of each fitted regression model for the four species in our analysis. The proportion variance explained is calculated as the difference between null and residual deviance divided by the null deviance. In silkworm, the mutation rate is modeled as a function of sequence context alone.

**S3 Table. Summary of CpG sites and SNPs for each species.** Genic regions are excluded in our analysis using NCBI *RefSeq* annotations. For human, phylogenetically conserved regions as defined by *phastcons* are also excluded. We identify C>T SNPs by polarizing the substitution that gave rise to each SNP using minor allele frequencies for human, chimpanzee, and rhesus macaque. For silkworm, we used *est-sfs*, a method that infers the ancestral state probabilities at polymorphic sites using information from a focal and outgroup species.

**S4 Table. Model comparison across CpG filtering and SNP polarization methods in human and non-human species.** Summary statistics of the 4-mer fitted regression model for each of the three primates in our analysis using the following combinations of CpG filtering and SNP polarization methods: intersected reference genome (CpGs present in both the hard-masked reference and inferred ancestral genome) and minor allele frequency (MAF) polarization; hard-masked reference genome and MAF polarization; intersected reference genome and ancestral allele polarization (ANC). The proportion variance explained is calculated as the difference between null and residual deviance, divided by the null deviance. We also show the number of CpG sites extracted from each genomic build after excluding genic regions and conserved regions (for human) and the number of CpG>TpG SNPs inferred using the corresponding polarization method.

## Notes

### Competing Interest Statement

The authors have declared no competing interest.

### Summary of Updates

Funder information added after Acknowledgements. No changes in analysis or results.

