## Supplementary Materials for "Sequence context and methylation interact to shape germline mutation rate variation at CpG sites"

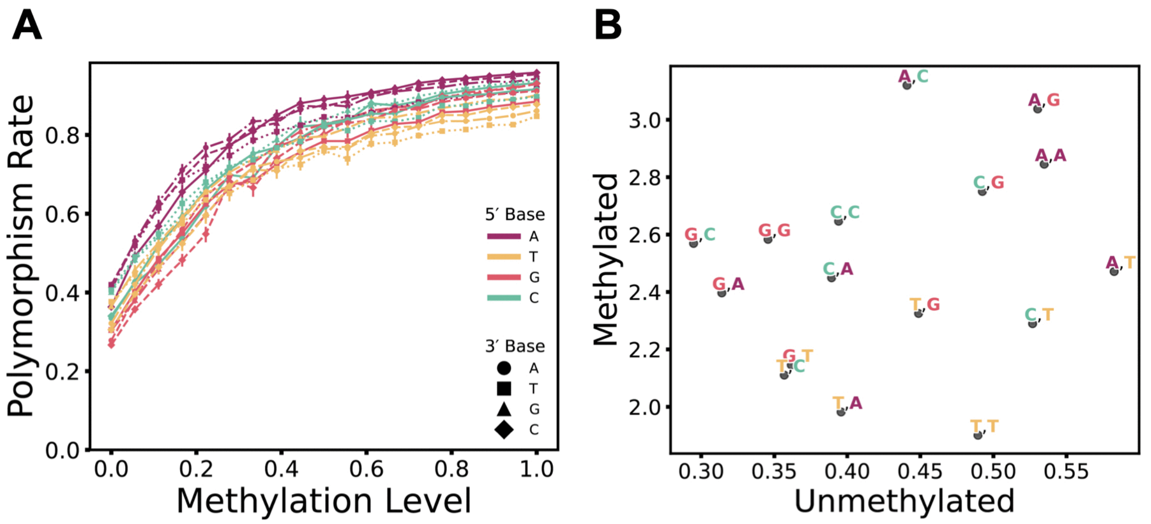


**S1 Fig. Interaction between the methyl group at the focal cytosine and flanking bases on CpG mutation rate. A.** Polymorphism rate for each 4-mer context, stratified by methylation levels. CpG sites were partitioned into 20 equal-width bins (ranging from [0, 0.05) to [0.95, 1]). Each point represents the proportion of SNPs among all CpG sites within a methylation bin, along with 95% confidence intervals assuming binomial sampling. Each context is represented by a color denoting the nucleotide at the 5' position and a shape denoting the nucleotide at the 3' position. **B.** Scaled mutation rates for each 4-mer context as estimated by the model for unmethylated and methylated sites. Points are labeled by the nucleotides upstream and downstream of the CpG, with error bars representing 95% confidence intervals for the estimates.

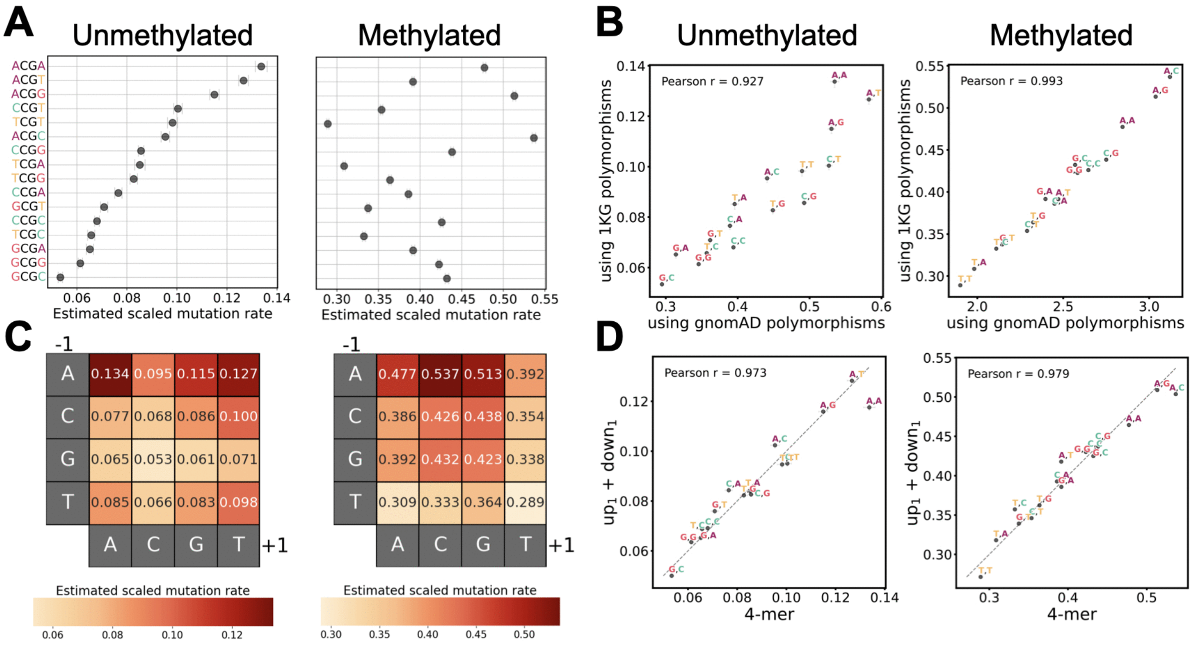


**S2 Fig. Concordance between mutation rates for 4-mer CpG contexts estimated from gnomAD (v4) and 1000 Genomes (1KG) polymorphism datasets. A.** Scaled mutation rate estimates for unmethylated and methylated sites for each 4-mer context using 1KG data. The contexts are sorted in descending order of mutability when unmethylated and error bars represent 95% confidence intervals for the estimates. **B.** Comparison of mutation rates estimated using polymorphisms from gnomAD and 1KG at unmethylated and methylated sites, respectively. **C.** Heatmap of scaled mutation rate estimates using the 4-mer model at unmethylated and methylated sites, using polymorphism data from 1KG. Each heatmap shows relative mutation rates for that panel; color scales are not comparable between panels. **D.** Concordance between mutation rates estimated from the 4-mer model and from the $up_1$+$down_1$ model using 1KG data. Points are labeled by the nucleotides upstream and downstream of the CpG, with error bars representing 95% confidence intervals for the estimates in panels B and D. The dashed line indicates $x = y$.

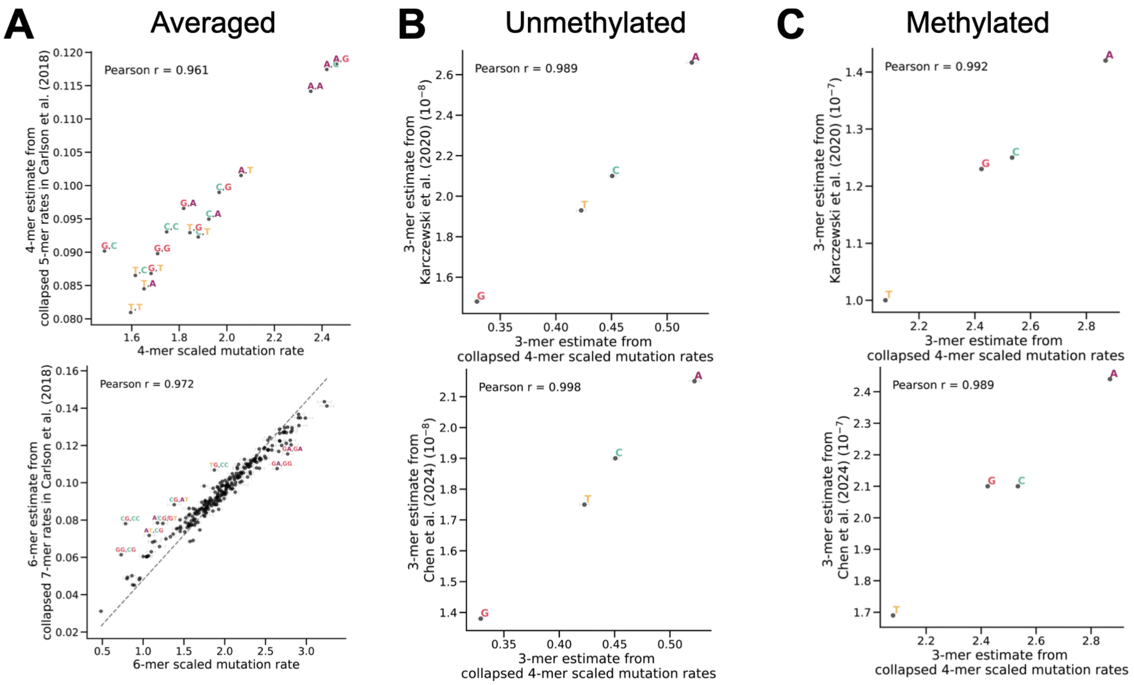


**S3 Fig. Context-specific scaled mutation rates are consistent with estimates from other published context-dependent mutation models. A.** Comparison with rates published in Carlson et al. (2018). To compare Carlson et al.'s estimates for 5-mers and 7-mers with our 4-mer and 6-mer results, we derived 4-mer and 6-mer rates as weighted averages of the mutation rate estimates for 5-mer (top) and 7-mer (bottom) contexts reported by Carlson et al., using the average methylation level of each 5-mer or 7-mer context as weights. For the expanded context comparison, a linear fit constrained through the origin is shown (dashed line). Only contexts with residuals exceeding 2.5 standard deviations from the fitted line are labeled. **B, C.** Comparison with rates published from gnomAD mutation rate models (Karczewski et al., 2020; Chen at al., 2024) for unmethylated (B) and methylated CpGs (C). To facilitate comparison, we computed the mutation rate for each 3-mer as the mean of estimates of the four corresponding 4-mer contexts sharing the same upstream base (e.g., rate for ACG is the mean of the rates of ACGA, ACGC, ACGG and ACGT). In all panels, points are labeled by the nucleotides upstream and downstream of the CpG, with horizontal error bars representing 95% confidence intervals for the model estimates.

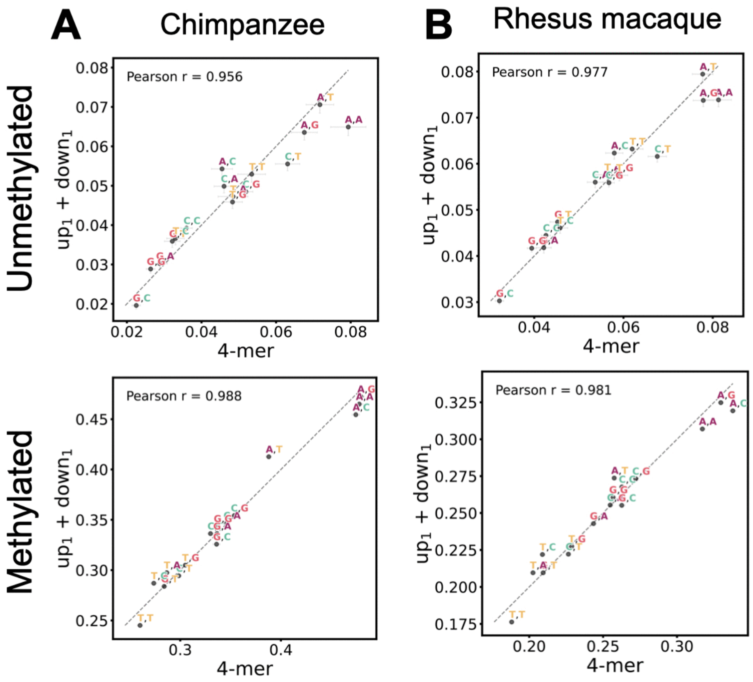


**S4 Fig. Independent effects of upstream and downstream bases on CpG mutability in chimpanzee and rhesus macaque. A.** Concordance between mutation rates as estimated from 4-mer model and from $up_1$ + $down_1$ model which assumes independent effects of upstream and downstream bases in chimpanzee for unmethylated and methylated sites. **B.** Same as (A), for rhesus macaque. Points are labeled by the upstream and downstream nucleotide flanking the CpG, and error bars denote 95% confidence intervals. The dashed line indicates $x = y$.

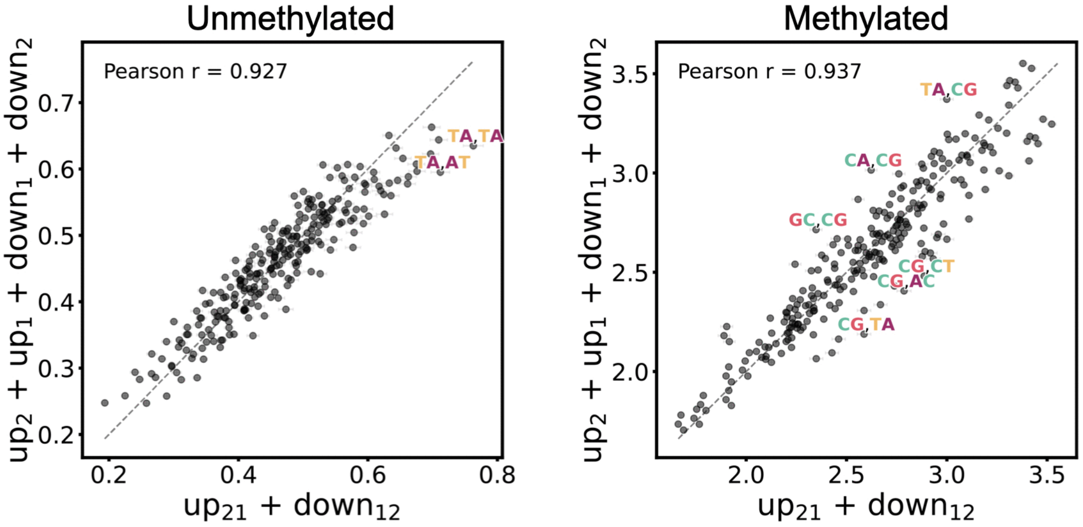


**S5 Fig. Concordance in mutation rate estimates between models with and without interactions within upstream and downstream dimers.** Comparison of human mutation rates as estimated from the $up_{21}$+$down_{12}$ model, which assumes interactions between bases within the dimer upstream and downstream to the CpG, and the $up_2$+$up_1$+$down_1$+$down_2$ model, which assumes independent effects of every flanking base. Only contexts for which the difference between the two models' estimates is more than 2.5 standard deviations away from the average difference across all contexts are labelled. Error bars represent 95% confidence intervals for the estimates on both axes. The dashed line indicates $x = y$.

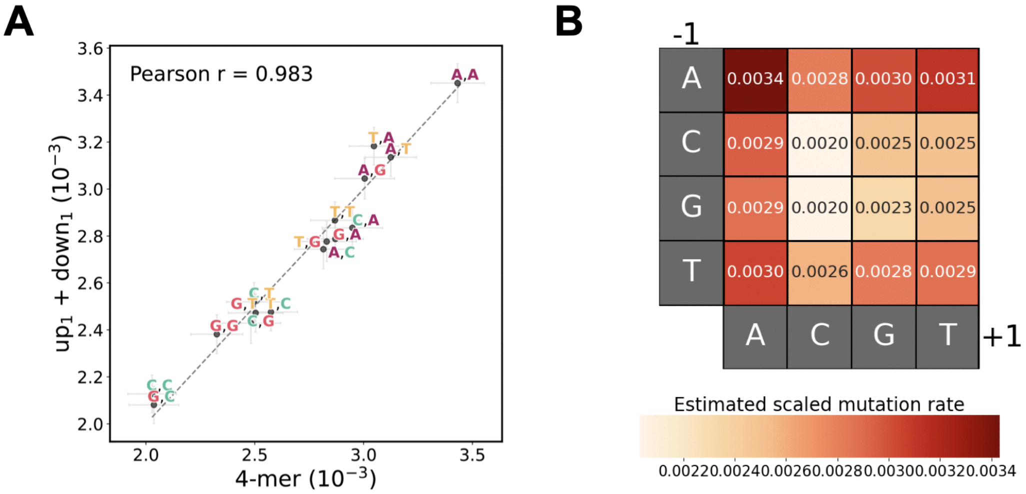


**S6 Fig. Independent effects of upstream and downstream bases on CpG mutability in the insect species, *Bombyx mori* (silkworm). A.** Concordance between mutation rates as estimated from 4-mer model and from $up_1$+$down_1$ model. Mutation rate is modelled as a function of sequence context alone**.** Points are labeled by the upstream and downstream nucleotide flanking the CpG, and error bars denote 95% confidence intervals. The dashed line indicates $x = y$. **B.** Heatmap of estimated scaled mutation rates using the 4-mer model.

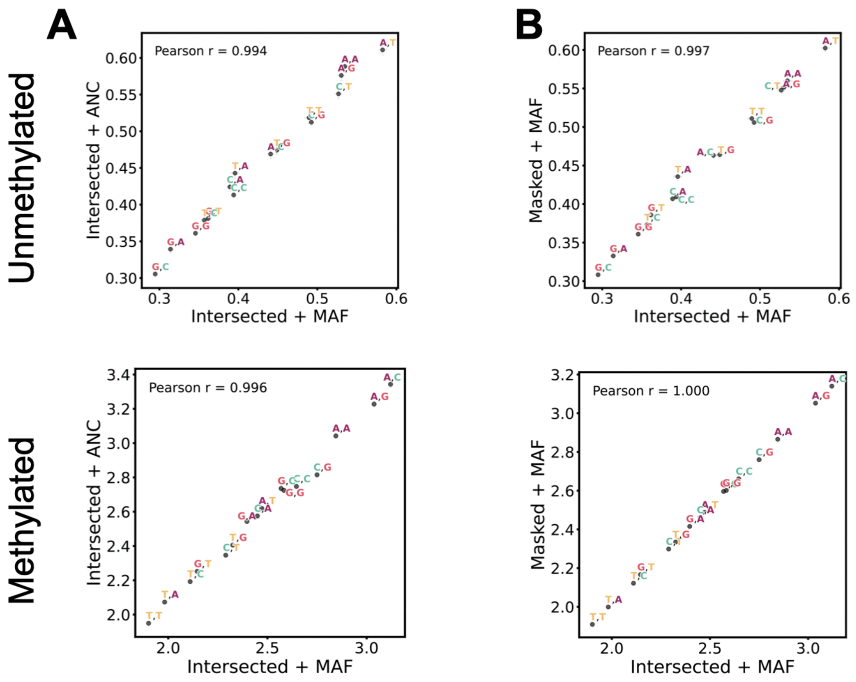


**S7 Fig. Context-specific scaled mutation rate estimates are highly concordant regardless of CpG filtering or SNP polarization method. A.** Estimates obtained by polarizing SNPs using minor allele frequency (MAF) are compared to those based on the inferred ancestral allele (ANC) for CpG sites present in both the hard-masked reference and inferred ancestral genome. **B.** Estimates derived from the intersected reference genome—restricted to CpGs present in both the hard-masked reference and inferred ancestral genome—are compared to those from the masked reference genome without intersection ("Masked"); SNPs are polarized by MAF. Points are labeled by the upstream and downstream nucleotide context flanking the CpG, and error bars denote 95% confidence intervals around the estimates.

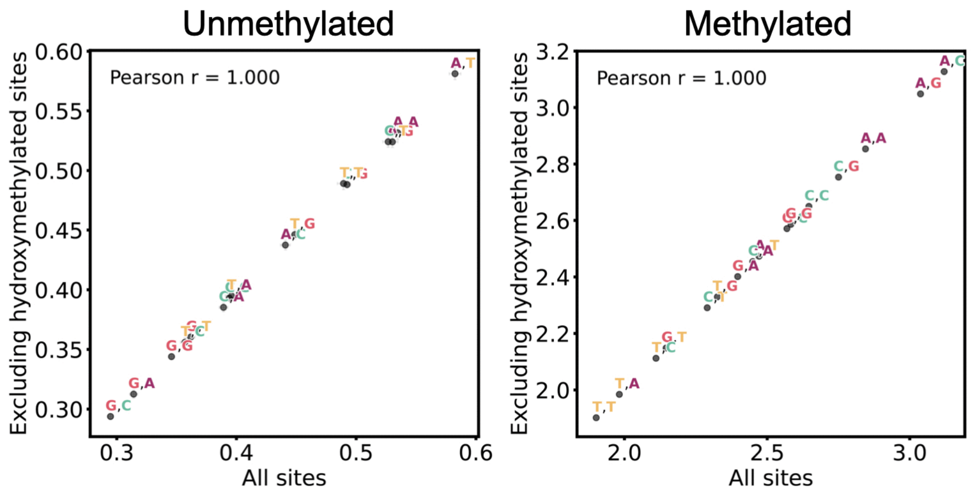


**S8 Fig**. **Scaled mutation rate estimates are nearly perfectly concordant after excluding hydroxy-methylated (5hmC) sites.** Comparison of the model's scaled mutation rate estimates for unmethylated and methylated sites when using all sites and when excluding sites with a non-zero 5hmC level as measured in human ES-cells. Points are labeled by the upstream and downstream nucleotide context flanking the CpG, and error bars denote 95% confidence intervals around the estimates.

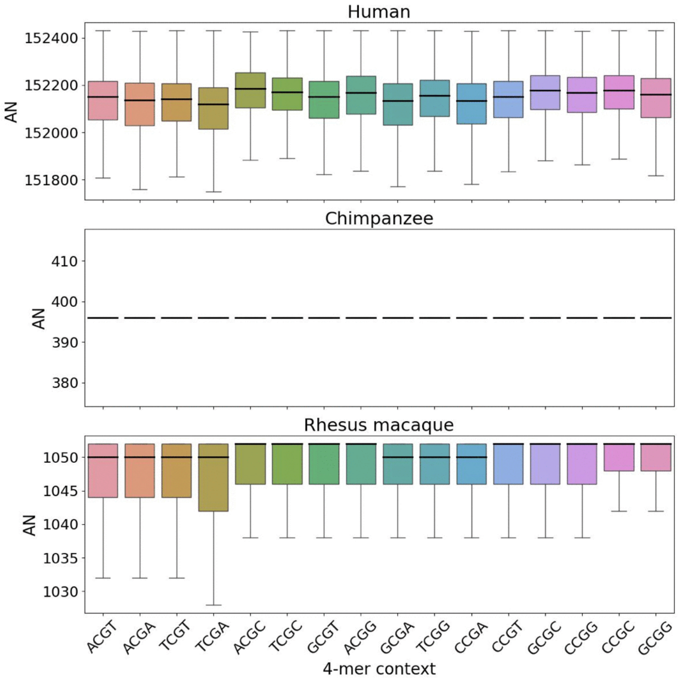


**S9 Fig. Callable allele coverage across 4-mer contexts in primate polymorphism datasets.** Distribution of the number of callable alleles by 4-mer sequence context in the human, chimpanzee, and rhesus macaque polymorphism dataset. Boxes show the median and interquartile range, with whiskers extending to 1.5x the interquartile range.

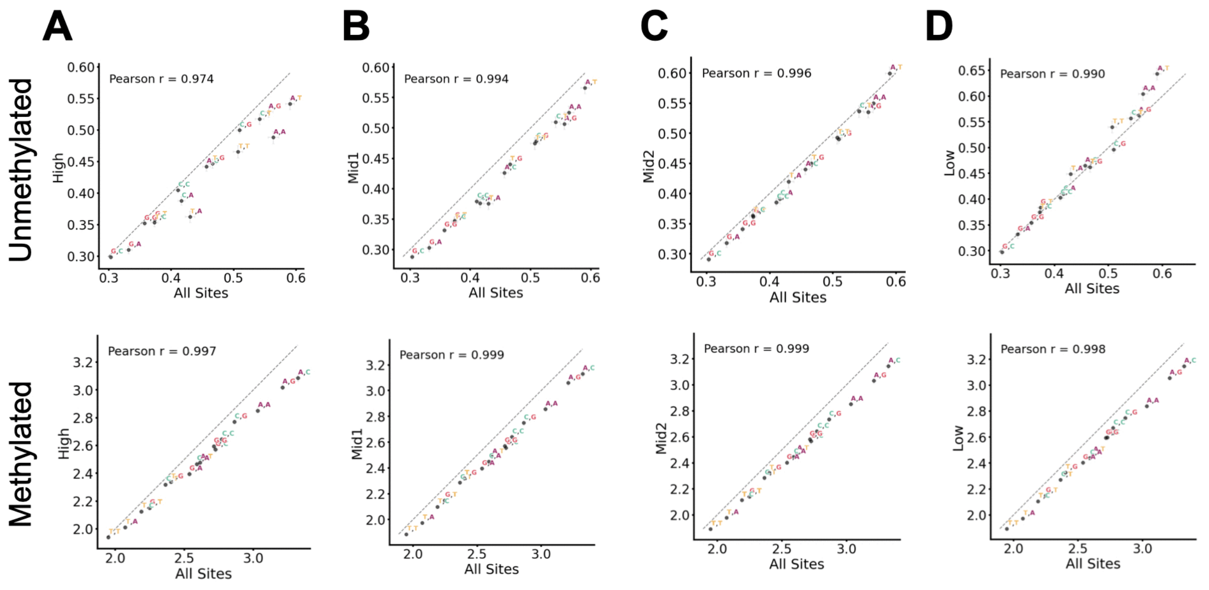


**S10 Fig. Context-specific scaled mutation rate estimates are consistent across recombination rate strata. A.** Comparison of scaled mutation rate estimates across all sites versus sites stratified by recombination rate, shown separately for unmethylated (top) and methylated (bottom) CpGs. Panels correspond to recombination rate quartiles: **A.** highest ("High", > 0.389), **B** upper-middle ("Mid1", ≤ 0.389), **C.** lower-middle ("Mid2", ≤ 0.026), and **D.** lowest ("Low", ≤ $4.877 \times 10^{-5}$). The dashed line indicates $x = y$. Points are labeled by the upstream and downstream nucleotide flanking the CpG, and error bars denote 95% confidence intervals.

**S1 Table. Scaled mutation rate estimates for each 4-mer and 6-mer CpG context at unmethylated and methylated sites (provided as excel file).** This table contains separate worksheets for each species and model (e.g., 4-mer, 6-mer), with each sheet reporting the sequence context, methylation status, dataset-specific scaled mutation rate estimate ($\mu T$), and its associated standard error. For silkworm, we report estimates from 4-mer and $up_1$+$down_1$ models only, and rates are modeled as a function of context alone.

| Species | Model | Null Deviance | Residual Deviance | Parameters | AIC | BIC | Proportion Variance Explained (%) |
|---|---|---|---|---|---|---|---|
| Human | 4-mer | 6,405,823 | 4,793,103 | 32 | 4,793,167 | 4,793,604 | 25.18% |
| | $up_1+down_1$ | 6,405,823 | 4,795,460 | 14 | 4,795,488 | 4,795,679 | 25.14% |
| | 6-mer | 6,405,823 | 4,771,028 | 512 | 4,772,052 | 4,779,036 | 25.52% |
| | $up_{21}+down_{12}$ | 6,405,823 | 4,776,600 | 62 | 4,776,724 | 4,777,569 | 25.43% |
| | $up_2+up_1+down_1+down_2$ | 6,405,823 | 4,783,251 | 26 | 4,783,303 | 4,783,658 | 25.33% |
| Chimpanzee | 4-mer | 4,464,404 | 4,235,923 | 32 | 4,235,987 | 4,236,410 | 5.12% |
| | $up_1+down_1$ | 4,464,404 | 4,237,096 | 14 | 4,237,124 | 4,237,309 | 5.09% |
| | 6-mer | 4,464,404 | 4,224,537 | 512 | 4,225,561 | 4,232,342 | 5.37% |
| | $up_{21}+down_{12}$ | 4,464,404 | 4,227,288 | 62 | 4,227,412 | 4,228,233 | 5.31% |
| | $up_2+up_1+down_1+down_2$ | 4,464,404 | 4,230,348 | 26 | 4,230,400 | 4,230,744 | 5.24% |
| Rhesus macaque | 4-mer | 8,809,643 | 8,516,789 | 32 | 8,516,853 | 8,517,303 | 3.32% |
| | $up_1+down_1$ | 8,809,643 | 8,518,893 | 14 | 8,518,921 | 8,519,118 | 3.30% |
| | 6-mer | 8,809,643 | 8,498,697 | 512 | 8,499,721 | 8,506,911 | 3.53% |
| | $up_{21}+down_{12}$ | 8,809,643 | 8,503,413 | 62 | 8,503,537 | 8,504,407 | 3.48% |
| | $up_2+up_1+down_1+down_2$ | 8,809,643 | 8,509,065 | 26 | 8,509,117 | 8,509,482 | 3.41% |
| Silkworm | 4-mer | 408,193 | 407,611 | 16 | 407,643 | 407,870 | 0.143% |
| | $up_1+down_1$ | 408,193 | 407,630 | 7 | 407,644 | 407,743 | 0.138% |

**S2 Table. Model comparison for human and non-human species.** Summary statistics of each fitted regression model for the four species in our analysis. The proportion variance explained is calculated as the difference between null and residual deviance divided by the null deviance. In silkworm, the mutation rate is modeled as a function of sequence context alone.

| | |
|---|---|
| **Human** | |
| Number of individuals | 807,162 |
| Number of non-genic, non-conserved CpG sites | 6,710,813 |
| Number of non-genic, non-conserved CpG C>T SNPs | 4,953,613 |
| **Chimpanzee** | |
| Number of individuals | 198 |
| Number of non-genic CpG sites | 8,523,730 |
| Number of non-genic CpG C>T SNPs | 958,654 |
| **Rhesus macaque** | |
| Number of individuals | 526 |
| Number of non-genic CpG sites | 9,505,961 |
| Number of non-genic CpG C>T SNPs | 1,714,326 |
| **Silkworm** | |
| Number of individuals | 1,082 |
| Number of non-genic CpG sites | 10,767,630 |
| Number of non-genic CpG C>T SNPs | 29,600 |

**S3 Table. Summary of CpG sites and SNPs for each species.** Genic regions are excluded in our analysis using NCBI *RefSeq* annotations. For human, phylogenetically conserved regions as defined by *phastcons* are also excluded. We identify C>T SNPs by polarizing the substitution that gave rise to each SNP using minor allele frequencies for human, chimpanzee, and rhesus macaque. For silkworm, we used *est-sfs*, a method that infers the ancestral state probabilities at polymorphic sites using information from a focal and outgroup species.

| Dataset | Null Deviance | Residual Deviance | Proportion Variance Explained (%) | Number of neutral sites | Number of CpG>TpG SNPs |
|---|---|---|---|---|---|
| Human (Intersected, MAF) | 6,405,823 | 4,793,103 | 25.18% | 6,710,813 | 4,953,613 |
| Human (Masked, MAF) | 7,418,266 | 5,635,875 | 24.03% | 8,176,931 | 5,908,808 |
| Human (Intersected, ANC) | 6,389,305 | 4,798,654 | 24.90% | 6,710,813 | 4,959,972 |
| Chimpanzee (Intersected, MAF) | 4,464,404 | 4,235,923 | 5.12% | 8,523,730 | 958,654 |
| Chimpanzee (Masked, MAF) | 4,866,736 | 4,616,709 | 5.14% | 10,010,745 | 1,046,806 |
| Chimpanzee (Intersected, ANC) | 4,294,376 | 4,083,915 | 4.90% | 8,523,730 | 891,262 |
| Rhesus macaque (Intersected, MAF) | 8,809,643 | 8,516,789 | 3.32% | 9,505,961 | 1,714,326 |
| Rhesus macaque (Masked, MAF) | 9,579,179 | 9,248,147 | 3.46% | 10,432,387 | 1,860,329 |
| Rhesus macaque (Intersected, ANC) | 6,925,334 | 6,761,980 | 2.36% | 9,505,961 | 1,158,913 |

**S4 Table. Model comparison across CpG filtering and SNP polarization methods in human and non-human species.** Summary statistics of the 4-mer fitted regression model for each of the three primates in our analysis using the following combinations of CpG filtering and SNP polarization methods: intersected reference genome (CpGs present in both the hard-masked reference and inferred ancestral genome) and minor allele frequency (MAF) polarization; hard-masked reference genome and MAF polarization; intersected reference genome and ancestral allele polarization (ANC). The proportion variance explained is calculated as the difference between null and residual deviance, divided by the null deviance. We also show the number of CpG sites extracted from each genomic build after excluding genic regions and conserved regions (for human) and the number of CpG>TpG SNPs inferred using the corresponding polarization method.
